# Mitochondrial dysfunction precedes neurodegeneration in DRPLA patient-derived neurons, and phenylbutyrate improves survival

**DOI:** 10.1101/2025.09.02.673807

**Authors:** Njoud Al Naama, Csaba Konrad, Shimaa Sherif, Christophe Raynaud, Chengbing Wang, Shenela Lakhani, Giovanni Manfredi, Caroline Alayne Pearson, M. Elizabeth Ross

## Abstract

Dentatorubral-pallidoluysian atrophy (DRPLA) is a progressive autosomal-dominant neurodegenerative disorder caused by a CAG repeat expansion in the ATN1 gene, which encodes the transcriptional corepressor Atrophin-1. The temporal sequence of molecular mechanisms driving neuronal dysfunction and degeneration in DRPLA is poorly understood, limiting therapeutic development. We generated patient-derived induced pluripotent stem cells and differentiated them into cortical excitatory glutamatergic neurons to model early pathogenic processes. Early pathological features of DRPLA patient-derived neurons included mitochondrial dysfunction and oxidative stress. These alterations occurred before overt neuronal loss, highlighting bioenergetic stress as a key early driver of disease progression toward neurodegeneration. Pharmacological treatment with phenylbutyrate significantly improved neuronal survival and reduced mitochondrial reactive oxygen species production, demonstrating the therapeutic potential of targeting mitochondrial dysfunction and oxidative stress. These findings challenge the conventional aggregation-centric model of polyglutamine disease pathogenesis and position mitochondrial stress as a central and early promoter of neuronal degeneration in DRPLA. By providing mechanistic insight into early-stage disease processes, our study lays the foundation for therapeutic strategies targeting mitochondrial dysfunction in DRPLA and related polyglutamine disorders.

## Introduction

Polyglutamine (PolyQ) expansion disorders like Huntington’s disease (HD) and spinocerebellar ataxias share key pathogenic mechanisms, including protein aggregation, disrupted proteostasis, and progressive neuronal degeneration (Tenchov *et al*, 2024). Among these PolyQ disorders, dentatorubral-pallidoluysian atrophy (DRPLA) remains one of the least understood mechanistically. DRPLA is associated with a CAG repeat expansion in the Atrophin 1 (ATN1) gene, which encodes a transcriptional corepressor. While trinucleotide expansion in ATN1 causes DRPLA, missense and insertion mutations in ATN1 cause a different syndrome of congenital hypotonia, epilepsy, profound developmental delay, digital anomalies (CHEDDA), and early death (Palmer *et al*, 2021; Wood *et al*, 2000; Zhang *et al*, 2006). The clinical manifestations of DRPLA vary based on the age of onset. Juvenile-onset cases typically present with intellectual disability and epilepsy, while adult-onset cases are characterized by ataxia, chorea, and dementia, progressive decline, and shortened life expectancy (Carroll *et al*, 2018). The molecular mechanisms underlying neurodegeneration in DRPLA, particularly those preceding overt neuronal loss, remain poorly defined. This knowledge gap has hindered the development of disease-modifying therapies, leaving DRPLA without effective treatments.

Existing DRPLA models have provided several insights into DRPLA pathology, but they fall short of capturing key features of the human disorder. Mouse models expressing expanded ATN1 develop nuclear inclusions, truncated protein fragments, autophagy dysfunction, motor deficits, and premature death (Baron *et al*, 2017; Schilling *et al*, 1999). However, surprisingly, these models do not exhibit significant neuronal loss, a hallmark of human DRPLA pathology. Postmortem studies provided valuable information on late-stage disease pathology (Takano *et al*, 1996; Yamada, 2010; Yamada *et al*, 2001; Yazawa *et al*, 1999), but they are limited in their ability to identify early pathogenic mechanisms. Cellular models, including patient-derived fibroblasts and mutant ATN1-expressing cell lines, have displayed accumulation of inclusions and autophagy defects (Baron *et al*., 2017; Suzuki *et al*, 2010). *Drosophila* models have also identified autophagy impairment and transcriptional dysregulation, with expression of mutant ATN1 leading to retinal degeneration and autophagosome accumulation linked to lysosomal clearance defects (Nisoli *et al*, 2010). Recent studies in flies highlighted dysregulation of protein quality control and immune response pathways as major contributors to DRPLA pathogenesis (Prifti *et al*, 2025). However, these models focus primarily on retinal degeneration, which is not present in DRPLA patients, limiting their translational relevance.

A key limitation in DRPLA research is the lack of human cell models that adequately recapitulate selective neuronal vulnerability and early-stage pathology. To address this gap, we leveraged patient-derived induced pluripotent stem cells (iPSCs) to model DRPLA pathology. In this study, we differentiated control and DRPLA patient-derived iPSCs into dorsal telencephalic neural progenitor cells (NPCs) and cortical excitatory neurons. Using this model, we systematically investigated early pathogenic events in DRPLA, including neuronal viability, protein aggregation, proteostatic dysfunction, oxidative stress, and mitochondrial impairment. Additionally, we evaluated the therapeutic potential of Sodium phenylbutyrate (PB) and Tauroursodeoxycholic acid (TUDCA), two compounds with established neuroprotective effects in HD, Alzheimer’s disease, and amyotrophic lateral sclerosis (Khalaf *et al*, 2022).

## Results

### DRPLA patient-derived iPSCs generate dorsal NPCs and cortical excitatory neurons with preserved early differentiation and network activity

Skin fibroblasts from four DRPLA patients and two healthy controls were reprogrammed into iPSCs. All cell lines were heterozygous for the ATN1 CAG expansion, carrying one non-pathogenic allele (<35 CAG repeats) and one pathogenic allele (> 60 CAG repeats) (EV Fig. 1A). Pluripotency was confirmed by immunostaining for canonical markers SSEA4 and SOX2 (Fig. 1A-B).

**Figure 1.**
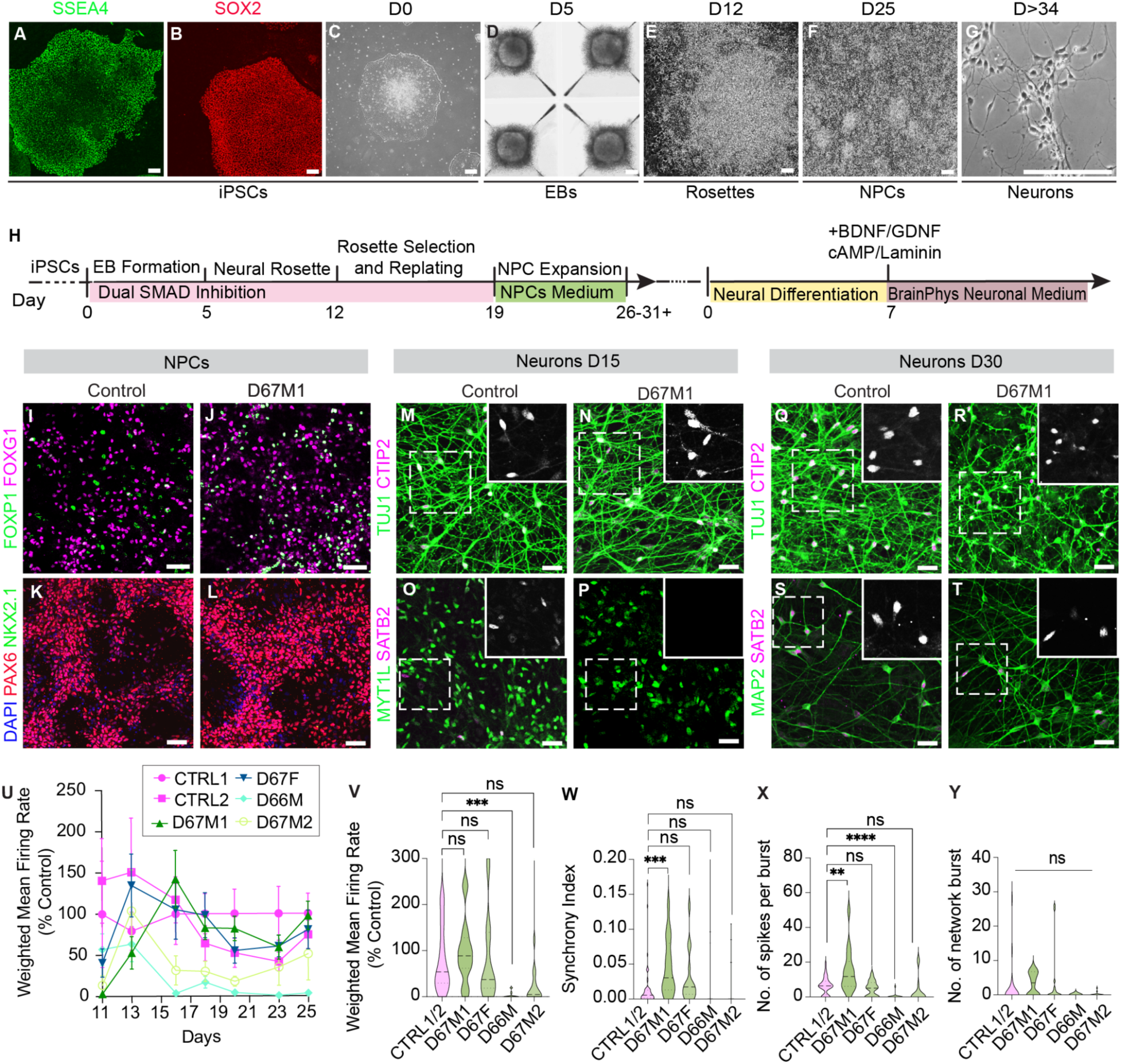
Generation and Characterization of control and DRPLA-Patient Derived Cortical NPCs and Neurons. (A-B) Immunostaining of iPSCs for pluripotency markers SSEA4 and SOX2. (C-G) Brightfield images of key stages of differentiation: iPSCs, embryoid bodies, neural rosettes, neural progenitor cells colonies, and cortical neurons, respectively. (H) Schematic overview of the cortical differentiation protocol. (I-J) Immunostaining for FOXP1 and FOXG1 in control and DRPLA patient-derived (D67M1) NPCs. (K-L) Immunostaining for PAX6, NKX2.1, and DAPI in control and DRPLA patient-derived (D67M1) NPCs. (M-N, Q-R) Immunostaining of TUJ1 and CTIP2 in control and patient-derived neurons at days 15 and 30. (O-P) Immunostaining of MYT1L and SATB2 in control and patient-derived neurons at days 15 and 30. (S-T) Immunostaining of MAP2 and SATB2 in control and patient-derived neurons at days 15 and 30. All insets show magenta channel only. (U) Weighted mean firing rate (% control) between differentiation day and 25. Data shown represents one experiment. N=3 technical replicates. (V-Y) Quantification of weighted mean firing rate (percent control), synchrony index, number of spikes per burst, and number of network bursts in control and DRPLA patient-derived neurons at day 25. Data are presented as mean ± SEM. N= 2 control and 4 patient-derived biological replicates, 3 technical replicates. p= ***0.0003 (V); ***0.0005 (W); **0.0015, ****0.0001(X). Statistical analysis was performed using unpaired Student’s t-test. Scale bars: (A-F) 100μM; (I-T) 50μM.

Using a well-established dual SMAD inhibition protocol (Chambers *et al*, 2009), iPSCs were directed toward a cortical lineage through defined developmental stages, including embryoid body formation, neural rosette emergence, and expansion of NPCs (Fig. 1C-H, EV Fig. 1B-D). NPCs expressed dorsal forebrain markers PAX6, FOXG1, and FOXP1, while lacking the ventral progenitor marker NKX2.1, confirming successful cortical lineage specification (Fig. 1I-L, EV Fig. 1E-J). Control and DRPLA patient-derived NPCs were subsequently differentiated into cortical excitatory neurons. Immunostaining revealed robust expression of TUJ1 (βIII-tubulin) and MAP2 (microtubule-associated protein 2), consistent with postmitotic neuronal identity. By day (D) 15 of differentiation, both the deep-layer projection neuron marker COUP-TF-interacting protein 2 (CTIP2) and Special AT-rich sequence-binding protein 2 (SATB2) were detected, with CTIP2 expression more prominent at early stages and SATB2 levels increasing by D30 (Fig. 1M-T, EV Fig. K-V). These findings indicate appropriate temporal subtype fate acquisition in control and DRPLA lines.

We performed multielectrode array (MEA) recordings to assess neuronal function at multiple differentiation time points. By D25, DRPLA neurons exhibited similar mean firing rates and network synchrony to controls, except for one line (D66M), which showed a reduced firing rate (Fig. 1U-W). Further analysis of network parameters, including the number of network bursts and spikes per burst, revealed no overt differences between genotypes (Fig. 1X-Y), suggesting that early-stage network activity is largely preserved in DRPLA patient-derived excitatory neurons. Together, these results demonstrate that DRPLA patient-derived iPSCs can be reproducibly differentiated into dorsal cortical NPCs and excitatory neurons with early-stage molecular and functional properties largely preserved, establishing a robust human *in vitro* system for investigating early-stage mechanisms of DRPLA pathogenesis.

### DRPLA patient-derived neurons exhibit early deep-layer fate bias and reduced upper-layer SATB2+ neuron generation

To determine whether DRPLA pathology impacted cortical fate specification, we quantified the percentage of MYT1L+ (*Myelin Transcription factor 1-like*) postmitotic neurons co-expressing either CTIP2 or SATB2 at D15, D30, and D45 of differentiation (Fig. 2A-L). At D15, DRPLA cultures showed a significantly higher proportion of CTIP2+/MYT1L+ neurons (∼83%) compared to controls (∼58%) (Fig. 2A, G, M), suggesting a premature commitment toward deep-layer cortical identity. By D30, CTIP2+ neuron proportions increased in control cultures to (∼72%), while DRPLA cultures declined to (∼58%) (Fig. 2C, I, M, EV Fig. 2A). From D30 to D45, CTIP2+ proportions remained stable within each genotype. However, by D45, DRPLA cultures exhibited a significantly lower percentage of CTIP2+ neurons (59%) compared to controls (∼75%) (Fig. 2E, K, M).

**Figure 2.**
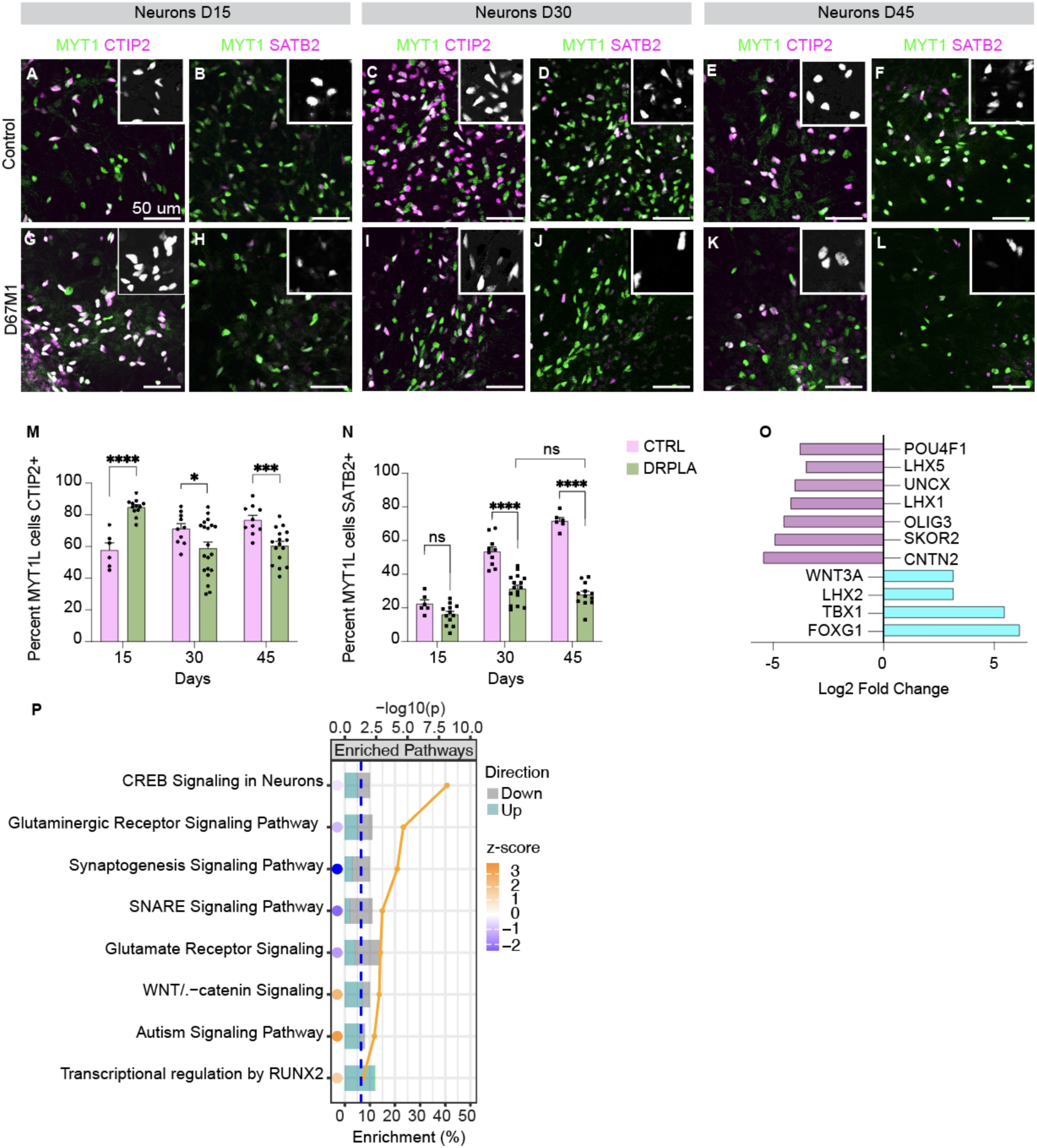
DRPLA patient-derived neurons exhibit an early shift toward deep-layer cortical fate differentiation. (A-L) Immunostaining of cortical fate markers CTIP2 and SATB2 co-labeled with the neuronal marker MYT1L in control and DRPLA patient-derived (D67M1) neurons at days 15, 30, and 45 of differentiation. (insets show magenta channel only). (M) Quantification of the proportion of MYT1L+ control and DRPLA patient-derived (D67M1) neurons expressing CTIP2. (N) Quantification of the proportion of MYT1L+ control and DRPLA patient-derived (D67M1) neurons expressing SATB2. (O) Misregulated neurogenesis genes identified in bulk RNA Seq analysis of control and DRPLA patient-derived neurons at day 20 of differentiation. (P) Gene ontological processes associated with neurogenesis were upregulated in DRPLA patient-derived neurons at day 20. Data are presented as mean ± SD. N=2 control and 4 patient-derived biological replicates, 3 technical replicates. Statistical analysis was performed using unpaired two-tailed Student’s t-tests. p= ****<0.0001, *0.0472, ***0.0007 (M); \*\*\*\**p*<0.0001 (N). Scale bars: (A-L) 50 μM.

At D15, there was no significant difference in the proportion of SATB2+/MYT1L+ neurons between control and DRPLA cultures (Fig. 2B, H, N). In control cultures, this proportion increased markedly from D15 to D30 and increased further between D30 and D45 (Fig. 2D, F, N). In contrast, DRPLA cultures showed a more limited increase between D15 and D30, with no significant change between D30 and D45, such that a significant reduction of SATB2+/MYT1L+ neurons were detected (Fig. 2H, J, L, N, EV Fig. 2B). This plateau suggests a reduced capacity of DRPLA neurons to generate or maintain upper-layer SATB2+ identity over time. These findings demonstrate that DRPLA patient-derived neurons exhibit a premature generation of deep-layer cortical neurons and a persistent reduction in upper-layer SATB2 neuron generation, highlighting altered temporal progression of cortical fate specification.

To identify molecular regulators that may underlie this altered fate trajectory, we performed bulk RNA sequencing on FACS-sorted pSyn-EGFP+ neurons derived from DRPLA patients and control iPSCs at D20 of differentiation. Gene expression analysis revealed significant dysregulation of several transcriptional factors involved in cortical development and neuronal subtype specification (Fig. 2O-P). Notably, FOXG1 has been shown to promote deep-layer fate CTIP2+, while delaying the emergence of upper layer neurons SATB2+ (Toma *et al*, 2014), was significantly upregulated in DRPLA neurons (Log2FC= +6.2) (Fig. 2P). Additional upregulated transcripts included *LHX2* and *WNT3A*, both of which have established roles in cortical progenitor patterning and excitatory neuron fate. Consistent with these gene-level changes, pathway enrichment analysis (IPA) revealed significant upregulation of WNT/β-catenin signaling, CREB signaling, as well as pathways related to synaptogenesis (Fig. 2O). These findings support the presence of premature differentiation in DRPLA neurons at the transcriptional level.

### Enhanced apoptosis and reduced survival in DRPLA patient-derived cortical excitatory neurons

To determine whether significant neurodegeneration occurred in DRPLA patient-derived neurons, we tracked over time after labeling postmitotic neurons using a pSyn-EGFP lentiviral reporter expressing EGFP under the neuron-specific human Synapsin promoter (Fig. 3A-D). Starting at D13 of differentiation, viable pSyn-EGFP+ neurons were quantified at regular intervals. DRPLA neurons showed a significant decline in survival as early as D20 (Fig. 3E), and by D70, all DRPLA lines showed a markedly reduced neuronal survival compared to controls (Fig. 3F).

**Figure 3.**
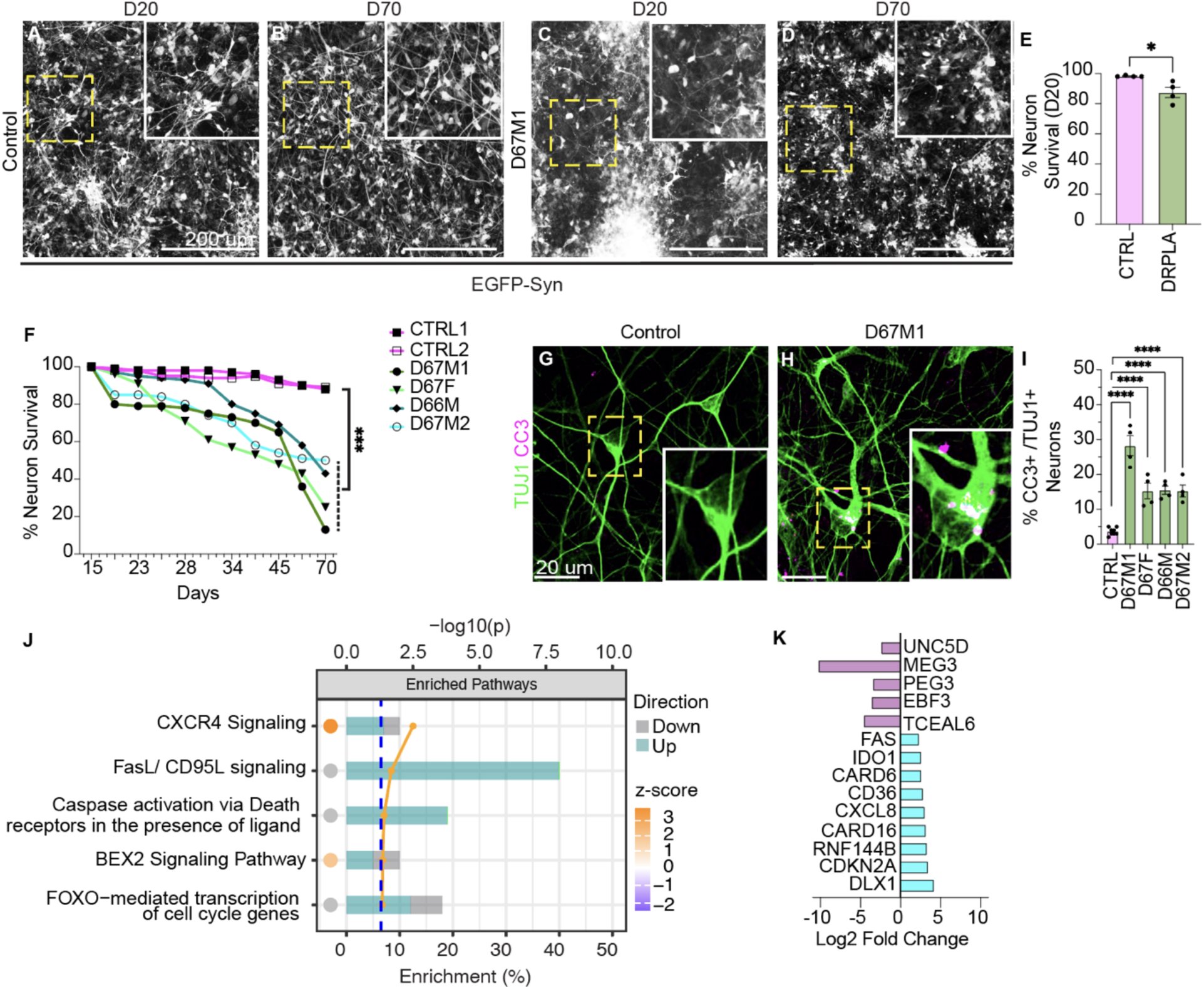
DRPLA patient-derived neurons exhibit increased apoptosis and progressive neuronal degeneration. (A-D) Representative images of pSyn-EGFP-labeled neurons in control and DRPLA patient-derived (D67M1) neurons at day 20 and day 70. Insets are high magnification images of boxed area. (E) Quantification of viable pSyn-GFP+ control and DRPLA patient-derived neurons at day 20. (F) Representative neuronal survival curve for controls and DRPLA patient-derived neurons from day 13 to 70. (G-H) Immunostaining of cleaved Caspase-3 (CC3) and TUJ1 in control and DRPLA (D67M1) patient-derived neurons at day 20. Insets show high magnification images of the boxed area. (I) Quantification of the percentage of TUJ1+ control and patient-derived neurons expressing CC3 at day 20. All data are presented as mean ± SEM. Statistical analysis was performed using unpaired Student’s t-test. N=2 control and 4 patient-derived biological replicates, 4 technical replicates. p= *0.0221 (E); ***0.0004 (F); ****< 0.0001 (I). Scale bars (A-D) 100 μM, (G-H) 20 μM.

Furthermore, we analyzed Cleaved Caspase 3 (CC3) levels in control and DRPLA patient-derived neurons. At D10, CC3 staining was undetectable in both groups (EV Fig. 2A-B). However, by D20, DRPLA patient-derived neurons exhibited a significantly increased proportion of CC3+ neurons relative to controls (Fig. 3G-I). At this stage, 20-30% of TUJ1+ DRPLA patient-derived neurons were CC3+, compared to fewer than 5% of control neurons (Fig. 3I), confirming elevated apoptotic activity in DRPLA patient-derived neurons. These results indicate that DRPLA patient-derived neurons undergo apoptotic cell death, as evidenced by increased CC3 levels and reduced long-term viability.

To identify transcriptional mechanisms contributing to this increased vulnerability, we interrogated our RNA-seq dataset from D20 neurons. Differential expression revealed enrichment of multiple apoptotic pathways, including FOXO-mediated transcription of pro-apoptotic genes, FasL/CD95 signaling, and caspase activation via death receptors (Fig. 3J). At the gene level, DRPLA neurons showed upregulation of several pro-apoptotic transcripts, including *FAS*, a canonical death receptor, as well as *IDO1*, *CARD6,* and *CARD16*, which are associated with inflammatory and caspase-dependent regulatory proteins (Karasawa *et al*, 2015; Sharma *et al*, 2019; Wang *et al*, 2019). In contrast, several genes involved in cell survival, such as *MEG3, UNC5D,* and *PEG3,* were downregulated, indicating possible loss of protective or compensatory transcriptional programs (Fig. 3K). Together, these data suggest that DRPLA patient-derived neurons exhibit a transcriptional profile consistent with apoptotic priming, characterized by concurrent activation of cell death pathways and downregulation of survival-promoting regulators.

### ATN1 inclusions are not detected in DRPLA patient-derived neurons under basal conditions

While nuclear inclusions of ATN1 are a hallmark of DRPLA postmortem pathology, it remains unclear whether aggregation occurs during early developmental stages or contributes to initial neuronal vulnerability. To address this, we assessed ATN1 expression and its aggregation propensity in DRPLA and control neurons across multiple time points. Immunocytochemistry (ICC) revealed a diffuse nuclear distribution of ATN1 in both groups, with no evidence of aggregate formation at D20 (Fig. 4A-E). Similarly, ICC for PolyQ-specific 1C2 antibody, which selectively detects expanded PolyQ repeats (>38 CAG repeats), did not detect PolyQ-positive inclusions in DRPLA patient-derived neurons at this stage (Fig. 4A-F). Western blot analysis revealed no difference in ATN1 protein level between control and DRPLA (EV Fig. 4A-B). We then performed a temporal analysis of ATN1 and PolyQ expression in differentiated neurons at D20, D40, and D90, confirming the absence of ATN1 and PolyQ-positive inclusions throughout neuronal maturation. (EV Fig. 4C-H). These data indicate that ATN1 aggregation is absent in DRPLA patient-derived neurons under basal conditions and is unlikely to be the primary driver accounting for neuronal cell death in this paradigm.

**Figure 4.**
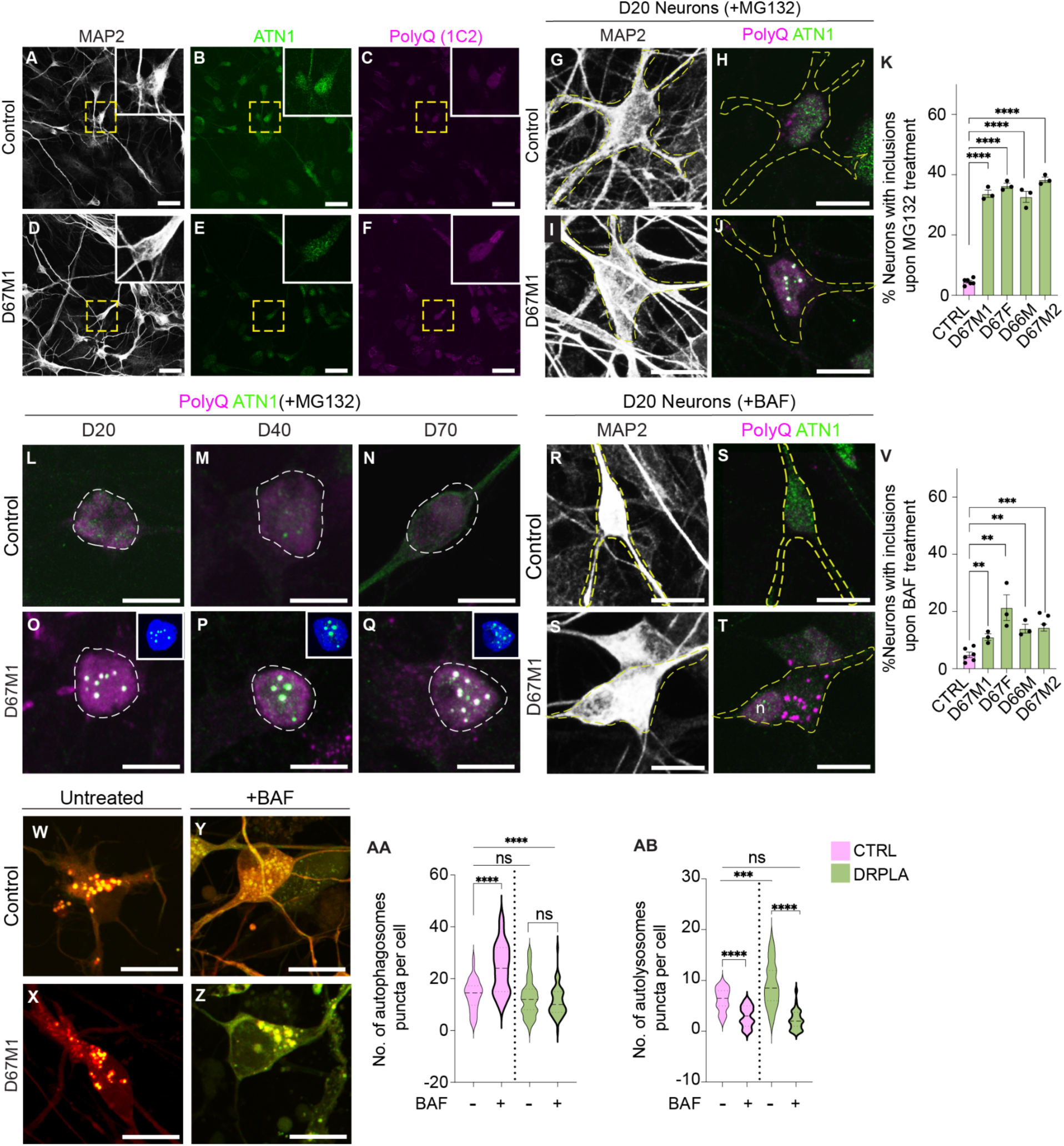
Absence of inclusions at basal levels and increased susceptibility to ATN1 inclusions in DRPLA patient-derived neurons with proteasome inhibition. (A-F) Immunostaining for MAP2, ATN1, and PolyQ (1C2) in control and DRPLA patient-derived neurons (D67M1) at day 20. (G-J) Immunostaining of MAP2, ATN1, and PolyQ in control and DRPLA patient-derived neurons treated with the proteasome inhibitor MG132 at day 20. (K) Quantification of the percentage of MAP2+ control and DRPLA patient-derived neurons treated with MG132 exhibiting inclusions at day 20. (L-Q) Immunostaining of ATN1 and PolyQ in control and DRPLA (D67M1) patient-derived neurons at days 20, 40, and 70. Insets in O-Q show DAPI+ nucleic localization of IBS. (R-T) Immunostaining of MAP2, PolyQ, and ATN1 in control and DRPLA patient-derived neurons following BAF application at day 20. n demarcates the position of the nucleus. (V) Quantification of the percentage of MAP2+ control and DRPLA patient-derived neurons with inclusions following BAF treatment. (W-Z) Representative images of untreated and BAF-treated control and DRPLA patient-derived (D67M1) neurons expressing the tandem monomeric mCherry-GFP-LC3 reporter. (AA-AB) Quantification of autophagosomes (mCherry+, GFP+ puncta) and autolysosomes (mCherry+, GFP-puncta) in control and DRPLA patient-derived neurons containing > 3 puncta per cell. N= 2 control and 4 patient-derived biological replicates, 3 technical replicates. Statistical analysis was performed using unpaired two-tailed Student’s t-tests. Data are presented as mean ± SD. p= ****<0.0001 (K); **0.0058 (D67M1), **0.0014 (D67F), **0.001 (D66M), ***0.0006 (D67F2) (V); ****<0.0001 (AA); ****<0.0001, ***0.0003 (AB). Scale bars (A-F) 20 μM (G-H) 10 μM (L-Q) 5 μM (R-T) 10 μM (W-Z) 10 μM.

### ATN1 aggregation is inducible upon proteostasis inhibition in DRPLA patient-derived neurons

To determine whether proteostasis mechanisms actively suppress ATN1 aggregation in DRPLA neurons, we pharmacologically inhibited the proteasome or autophagy in DRPLA and control neurons. At D20 of neuronal differentiation, treatment with the proteasome inhibitor MG132 (5 µM, 18h) selectively induced intranuclear ATN1 inclusions in DRPLA neurons but not controls, as confirmed by the co-localization of ATN1 and PolyQ antibodies (Fig. 4G-K). We then asked whether aged neurons had a higher propensity to form inclusions when challenged by comparing ATN1 aggregation following proteosome inhibition at D20, D40, and D70. However, the proportion of neurons harboring intranuclear ATN1 inclusions remained unchanged across each time point (Fig. 4L-Q). This suggests the age of the neuron is not a factor in the propensity of DRPLA patient-derived neurons to form inclusions upon inhibition of proteosomal pathways.

To evaluate the role of the autophagy-lysosomal pathway in aggregate clearance, we treated D20 neurons with Bafilomycin A1 (BAF) (100 nM, 18h) to block autophagosome-lysosome fusion. This led to cytoplasmic (not nuclear) PolyQ-positive inclusions in DRPLA patient-derived neurons, indicating that autophagy inhibition triggers a distinct subcellular pattern of aggregation (Fig. 4R-V). To further investigate autophagy dynamics, we utilized the mCherry-GFP-LC3 reporter to distinguish between autophagosomes (mCherry+GFP+) and autolysosomes (mCherry+GFP-). Under basal conditions, DRPLA patient-derived neurons exhibited a significant accumulation of autolysosomes compared to controls (Fig. 4W, X), while autophagosome levels were comparable between groups (Fig. 4AA-AB). Following BAF treatment, control neurons showed the expected increase in autophagosomes, whereas DRPLA patient-derived neurons failed to respond (Fig. 4Y, Z, AA), despite maintaining autolysosome numbers comparable to controls (Fig. 4AB). This suggests that DRPLA patient-derived neurons accumulate autolysosomes at baseline and exhibit impaired autophagosome biogenesis or turnover when challenged. These findings indicate that ATN1 aggregation is not an intrinsic feature of DRPLA neurons under basal conditions. However, the appearance of ATN1 inclusions following proteasome or autophagy inhibition indicates that DRPLA neurons depend on protein quality control systems to prevent ATN1 aggregation.

### Mitochondrial hyperpolarization, elevated oxidative stress, and altered morphology in DRPLA patient-derived neurons

Since no detectable ATN1 inclusions were observed in DRPLA patient-derived neurons at baseline, we asked whether elevated oxidative stress contributes to decreased neuronal survival. We first assessed accumulation of reactive oxygen species (ROS) using live dihydroethidium (DHE) imaging in pSyn-EGFP+ neurons. Quantification of fluorescence intensity over time revealed a significant increase in ROS production in DRPLA lines (Fig. 5A-E). At D20, DRPLA patient-derived neurons exhibited an approximate 150% increase in the accumulation of DHE fluorescence intensity over 20 minutes compared to controls (Fig. 5A-E). The mitochondrial contribution to ROS generation was examined using MitoSOX, a fluorescent indicator of mitochondrial ROS (mtROS). DRPLA patient-derived neurons exhibited a 2- to 3-fold increase in mtROS compared to controls (Fig. 5F-J), suggesting mitochondria are a key source of elevated oxidative stress.

**Figure 5.**
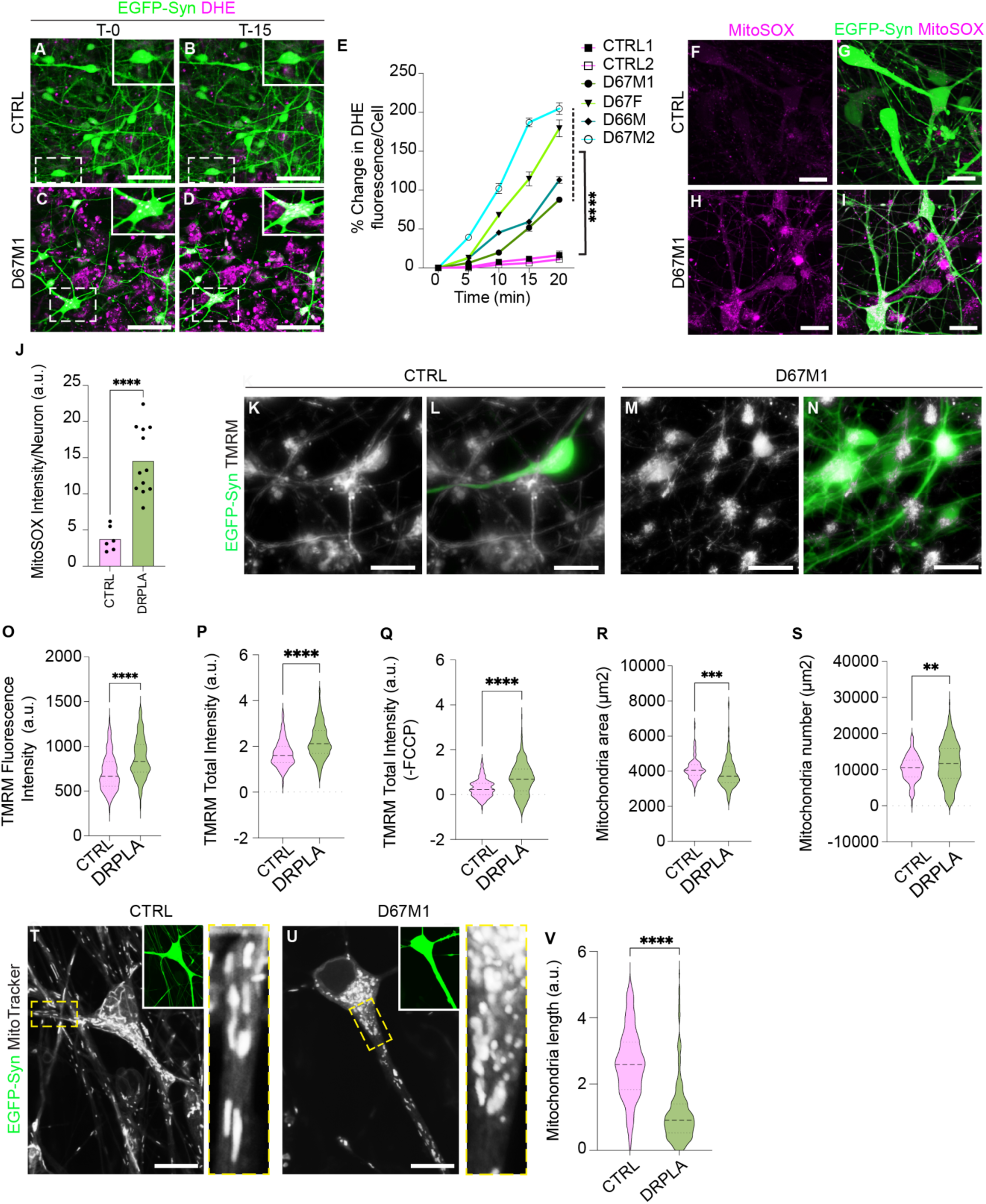
DRPLA patient-derived neurons exhibit elevated oxidative stress and dysfunctional mitochondria. (A-D) Representative live-cell images of control and DRPLA (D67M1) patient-derived neurons expressing pSyn-EGFP and treated with dihydroethidium (DHE) at baseline (T-0) and after 15 minutes (T-15). Insets show magnified views of individual neuronal somas. (E) Quantification of the percent change in DHE fluorescence intensity per neuron over time in control and DRPLA patient-derived neurons. Each point represents the average of DHE fluorescence intensity from all neurons, pooled from three independent differentiations. (F-I) Representative images showing MitoSOX staining in pSyn-EGFP-expressing control and DRPLA patient-derived neurons at day 20. (J) Quantification of MitoSOX fluorescence intensity in control and DRPLA patient-derived neurons at day 20. (K-N) Representative live-cell images of control and DRPLA patient-derived (D67M1) neurons at day 20 of differentiation, expressing pSyn-EGFP and stained with TMRM (ΔΨm). (O-S) Quantification of mitochondrial TMRM fluorescence intensity, total TMRM signal, and FCCP-subtracted TMRM signal, area, and number in control and DRPLA patient-derived neurons at day 20. (T-U) Representative images of mitochondria labeled with MitoTracker in control and DRPLA patient-derived neurons at day 20. Insets show high magnification images demarcated by yellow dashed boxes. (V) Quantification of mitochondrial length in control and DRPLA patient-derived neurons at day 20. Data represent mean ± SEM. Statistical analysis was performed using unpaired two-tailed Student’s *t*-test. N=2 control and 4 patient-derived biological replicates, 3 technical replicates. p=****<0.0001 (E); ****<0.0001 (J); ****<0.0001 (O-Q); ***0.0001 (R); **0.003. (V) *P* < 0.0001. Scale bars: (A-B) 50 μM, (F-N) 20 μM, (T-U) 10μM.

Transcriptomic data further supported these findings. Differential gene expression analysis revealed downregulation of glutathione metabolism genes, including *GSTM1* and *GSTM5*. GSTM enzymes are detoxifying and use glutathione (GSH). The combination of elevated ROS generation at the same time GSTM expression is decreased suggests an impaired antioxidant defense capacity in DRPLA patient-derived neurons (EV Fig. 5G). In contrast, pro-inflammatory mediators, including *CCL2*, *IL32,* and *PSMB8/9* were significantly upregulated, indicating activation of inflammatory cascades and potential ROS amplification. Additional genes linked to mitochondrial function and stress response, such as *PDK4, GATD3, LRRK2*, and *HSPB6,* were upregulated. IPA enrichment analysis revealed activation of mitochondrial division signaling, NRF2 (NFE2L2)-mediated oxidative stress responses, iron homeostasis, and energy metabolism (EV Fig. 5F). These transcriptomic signatures support a model in which mitochondrial dysfunction and redox imbalance are early, intrinsic features of DRPLA pathology.

We next evaluated mitochondrial bioenergetics to assess mitochondrial membrane potential (ΔΨm), as it correlates to mtROS (Suzuki, 2012), using live-cell tetramethylrhodamine methyl ester (TMRM) fluorescence in D20 pSyn-EGFP+ neurons (Fig. 5K-N). Under basal conditions, DRPLA patient-derived neurons showed significantly elevated whole neuron TMRM mean fluorescence intensity compared to controls, indicating increased mitochondrial ΔΨm per unit area (hyperpolarization) (Fig. 5O). In parallel, total TMRM intensity, which integrates both ΔΨm and mitochondrial content, was also markedly elevated in DRPLA neurons (Fig. 5P), and upon carbonyl cyanide-p-trifluoromethoxyphenylhydrazone (FCCP, 100 μM) treatment, an uncoupling agent that dissipates ΔΨm, both groups showed a reduction in total TMRM signal, but DRPLA neurons retained significantly higher TMRM levels (Fig. 5Q), suggesting increased resistance to depolarization.

In view of the significant increase in neuronal death at D20 (Fig. 3H), we next asked whether mitochondrial hyperpolarization is an early event preceding neuronal loss. To address this, we assessed ΔΨm at D10, a stage before widespread neuronal loss, and observed a significant elevation in mitochondrial ΔΨm (EV Fig. 5A). Total TMRM fluorescence intensity at baseline and following FCCP treatment was also significantly elevated at D10 (EV Fig. 5B-C), indicating that mitochondrial hyperpolarization is an early feature of the disease.

We analyzed mitochondrial morphology using TMRM and MitoTracker staining to further assess mitochondrial integrity. TMRM analysis revealed a significant reduction in mitochondrial area in DRPLA patient-derived neurons at both D20 and D10 (Fig. 5R, EV Fig. 5D). Mitochondrial number per neuron was significantly higher in DRPLA patient-derived neurons at D20, but not D10 (Fig. 5S, EV Fig. 5E), suggesting fragmentation of the mitochondrial network at D20.

MitoTracker staining confirmed a predominance of small, rounded mitochondria in DRPLA patient-derived neurons, in contrast to the elongated interconnected mitochondrial network observed in control neurons (Fig. 5T-V). These findings reveal early and sustained mitochondrial dysfunction, marked by hyperpolarization, altered morphology, and elevated oxidative stress in DRPLA patient-derived neurons. This early bioenergetic dysregulation emerges before neuronal loss, suggesting it is a primary pathogenic mechanism rather than a downstream consequence of degeneration.

### PB alone is sufficient to reduce oxidative stress and increase survival in DRPLA patient-derived neurons

We next asked whether pharmacological intervention could mitigate neuronal degeneration, selecting sodium phenylbutyrate (PB) and tauroursodeoxycholic acid (TUDCA) as proof-of-principle compounds based on their reported neuroprotective properties (Fels *et al*, 2022; Romero-Ramirez & Mey, 2024). Their therapeutic potential in DRPLA was systematically assessed as monotherapies and in combination across a range of concentrations. PB was tested at 1000 µM, 500 µM, and 100 µM, TUDCA at 100 µM, 50 µM, and 25 µM. In addition, five PB-TUDCA combinations were assessed (1000µM PB-100 µM TUDCA; 1000µM PB-50 µM TUDCA; 500µM PB-50 µM TUDCA; 500µM PB-25 µM TUDCA; 100µM PB-10 µM TUDCA). Treatments were initiated on D1 of neuronal differentiation, administered daily through D7, and maintained thereafter in maturation media.

#### Effect on Oxidative Stress

Given the elevated oxidative stress observed in DRPLA neurons, we first tested whether pharmacological treatment with PB alone or with TUDCA could attenuate ROS accumulation. Untreated DRPLA patient-derived neurons exhibited significantly increased DHE intensity compared to controls, indicating elevated oxidative stress (Fig. 6A-D, M). PB (1000 µM) was selected based on prior reports supporting its efficacy across diverse experimental models (Braun *et al*, 2022; Fels *et al*., 2022; Hu *et al*, 2025; Zhang *et al*, 2025). Strikingly, treatment with PB (1000 µM) and co-treatment with TUDCA (100 µM) reduced DHE signal and accumulation to control levels (Fig. 6E-L, N, O). DRPLA neurons also showed a significantly elevated MitoSOX fluorescence relative to controls (Fig. 6P-S, AB), and treatment with PB or PB-TUDCA significantly diminished mtROS in DRPLA patient-derived neurons to control levels (Fig. 6T-AA, AC-AD). These results demonstrate that application of PB or PB-TUDCA effectively attenuates ROS in DRPLA patient-derived neurons.

**Figure 6.**
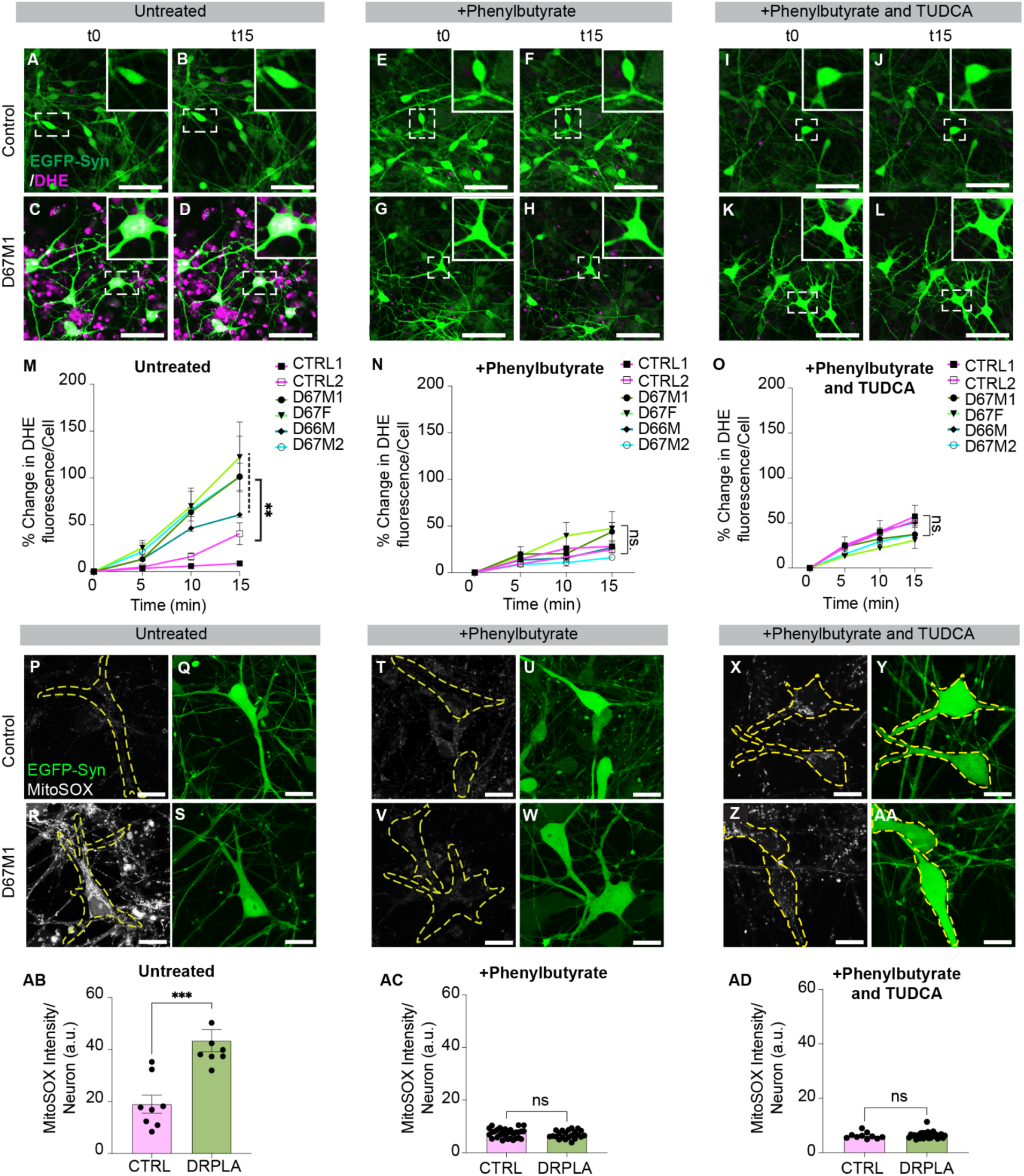
Phenylbutyrate monotherapy and Phenylbutyrate-TUDCA combination reduces cytosolic and mtROS in DRPLA patient-derived neurons. (A-D) Representative images of DHE staining in untreated day 20 pSyn-EGFP-labeled control and DRPLA patient-derived neurons at time point 0 (t0) and after 15 minutes (t15). (E-H) Representative images of DHE staining in day 20 pSyn-EGFP-labeled control and DRPLA patient-derived neurons treated with 1000 µM phenylbutyrate at t0 and t15. (I-L) Representative images of DHE staining in day 20 pSyn-EGFP-labeled control and DRPLA patient-derived neurons treated with 1000 µM phenylbutyrate and 100µM at t0 and t15. Insets show magnified views of DHE-positive neurons. (M) Quantification of DHE fluorescence over time in untreated control and DRPLA patient-derived neurons, shown as percent change in fluorescence per neuron relative to baseline (t0). (N) Quantification of DHE fluorescence over time in 1000 µM phenylbutyrate treated control and DRPLA patient-derived neurons, shown as percent change in fluorescence per neuron relative to baseline (t0). (O) Quantification of DHE fluorescence over time in 1000 µM phenylbutyrate/100 µM TUDCA treated control and DRPLA patient-derived neurons, shown as percent change in fluorescence per neuron relative to baseline (t0). (P-S) Representative images of mitochondrial ROS detected by MitoSOX in untreated control and DRPLA patient-derived neurons at day 20. (T-W) Representative images of mitochondrial ROS detected by MitoSOX in 1000 µM phenylbutyrate treated control and DRPLA patient-derived neurons at day 20. (X-AA) Representative images of mitochondrial ROS detected by MitoSOX in 1000 µM phenylbutyrate/100 µM TUDCA treated control and DRPLA patient-derived neurons at day 20. (AB-AD) Quantification of MitoSOX intensity in untreated, 1000 µM phenylbutyrate, and 1000 µM phenylbutyrate/100 µM TUDCA-treated pSyn-GFP+ control and DRPLA patient-derived neurons at day 20. Yellow dashed outlines indicate neuronal somata. Data represent mean ± SEM. Statistical analyses were performed using a two-tailed Student’s *t*-test. N=2 control and 4 patient-derived biological replicates, 3 technical replicates. p=**0.0011 (M); *** 0.0006 (AB). Scale bars: (A-L) 50 μM (P-AA) 20 μM.

#### Effect on neuronal survival

Having established that these compounds reduce oxidative stress, we next sought the optimal treatments and doses to promote long-term neuronal survival. We performed longitudinal survival assays on control and DRPLA patient-derived EGFP-Syn+ neurons over 70 days to determine whether PB alone improved neuronal survival. In the absence of treatment, DRPLA patient-derived neurons exhibited progressive neuronal loss without treatment, with a steep decline observed after D20 (Fig. 7A-D, M). PB treatment significantly increased survival in all DRPLA lines, with PB 1000µM producing the most robust and consistent rescue across lines (Fig. 7E-H, N, Q, EV Fig. 6A). PB 500µM significantly improved survival at D40 (EV Fig. 5A, EV Fig. 6A), while PB 100µM enhanced survival in three DRPLA lines but reduced survival in D67M2 compared to untreated (EV Fig. 5B, EV Fig. 6A). By day 70, survival in untreated DRPLA patient-derived neurons dropped below 40%, whereas controls maintained stable levels (EV Fig. 6D). PB 1000µM continued to significantly increase survival across all patient lines (EV Fig. 6D). In contrast, PB 500µM and 100µM showed a significant rescue in only one patient line, but no benefit in the others (EV Fig. 6D). Notably, PB treatment reduced survival in control neurons at all concentrations (Fig. 8E-F, N, P, EV Fig. 6D), suggesting a disease-specific effect. Comparison of D40 and D70 data indicates that PB 1000µM provided a robust and sustained neuroprotective effect across DRPLA lines (EV Fig. 6A, D), suggesting a lasting neuroprotective benefit when administered at higher concentrations.

**Figure 7.**
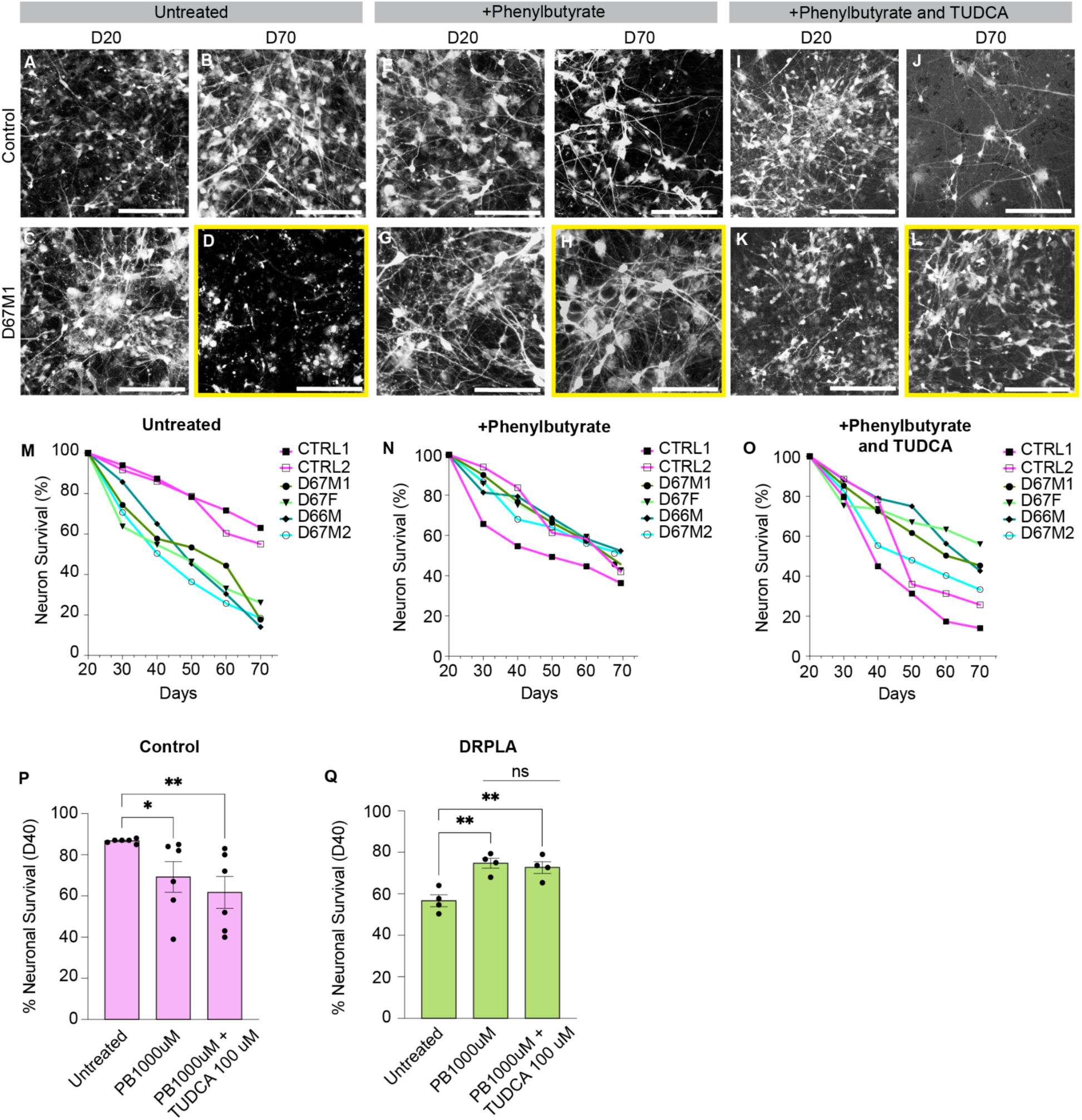
Phenylbutyrate monotherapy rescues neuronal survival in DRPLA patient-derived neurons. (A-D) Representative images of untreated pSyn-EGFP-labeled control and patient-derived neurons on days 20 and 70 of differentiation. (E-H) Representative images of 1000µM Phenylbutyrate treated pSyn-EGFP-labeled control and patient-derived neurons days 20 and 70 of differentiation. (I-L) Representative images of 1000µM Phenylbutyrate/100µM TUDCA (determined optimal concentrations) treated pSyn-EGFP-labeled control and patient-derived neurons days 20 and 70 of differentiation. Yellow boxed images represent end endpoint for comparison across untreated and treated DRPLA patient-derived neurons. (M-O) Longitudinal survival curves for each control and DRPLA patient-derived cell line under the three treatment conditions. Neuronal survival was quantified as the percentage of EGFP+ neurons at each time point normalized to the first day of imaging. Each curve represents the mean survival from three independent differentiations per line. (P) Quantification of neuronal survival in control neurons at day 40. Data represent mean ± SEM. N=2 control and 4 patient-derived biological replicates, 3 technical replicates. Statistical analysis was performed using two-tailed Student’s t-test. p = *0.0415, **0.0093 (P); **p=0.0030, **p=0.0073 (Q). Scale bar: (A-L) 50 μM.

The ability of TUDCA monotherapy to provide neuroprotection was assessed. At D40, TUDCA 100µM significantly improved survival in all DRPLA lines except D67M2. (EV Fig. 5C, EV Fig. 6B). However, by D70, this protective effect was sustained only in D67M1, with no significant improvement observed in the other lines (EV Fig. 6E). Lower concentrations of TUDCA (50-10µM) did not produce significant improvements at either time point (EV Fig. 5D-F, EV Fig. 6 B, E). As with PB, TUDCA treatment reduced survival in control neurons, indicating possible toxicity in non-diseased cells (EV Fig. 6 B, E). These results suggest that TUDCA provides early but limited neuroprotection, with 100µM being the most effective dose, but lacking efficacy over time.

We asked whether combining PB and TUDCA could further enhance neuroprotection. At D40, the highest-dose combination (1000µM PB + 100µM TUDCA) significantly increased neuronal survival in all DRPLA lines except D66M (Fig. 7I-L, O, Q); however, by D70, this combination provided significant rescue across all DRPLA lines (EV Fig. 6E). Intermediate combinations such as 1000µM PB + 50µM TUDCA showed moderate benefits in select DRPLA lines at D40 but were largely ineffective at D70 (EV Fig. 5G, EV Fig. 6C, F). Similarly, 500µM PB + 50µM TUDCA and 500µM PB + 25 µM TUDCA significantly improved survival in some lines at D40 but failed to sustain protection at D70 (EV Fig. 5H-I, EV Fig. 6C, F). The lowest-dose combination 100µM PB + 10µM TUDCA did not enhance survival at either time point (EV Fig. 5J, EV Fig. 6C and F). Overall, the combination treatment exhibited variable effects across patient lines, highlighting the heterogeneity in treatment sensitivity among DRPLA-derived neurons. The survival levels with the combination treatment did not outperform those observed with PB 1000µM alone. Both PB treatment alone and in combination with TUDCA exhibited adverse effects on wild-type neurons compared to untreated wild-type neurons (Figure 7M-Q). In summary, both PB and the combination therapy with TUDCA effectively reduced neuronal cell death in DRPLA patient-derived neurons (Figure 7Q). However, chronic treatment with 1000µM PB was the most effective, as evidenced by its significant reduction of oxidative stress and neuronal survival.

## Discussion

This study uncovers early features of cellular pathogenesis in DRPLA by leveraging patient-derived iPSCs to model disease onset. While PolyQ disorders have traditionally been attributed to aggregate-mediated toxicity, our findings reveal that mitochondrial hyperpolarization and oxidative stress precede neuronal loss and ATN1 inclusions.

Mitochondrial hyperpolarization was detected as early as D10 of neuronal differentiation, prior to a detectable increase in ROS and mitochondrial fragmentation, suggesting a bioenergetic imbalance intrinsic to DRPLA. These bioenergetic alterations could be the result of an imbalance between electron flow in the respiratory chain and ATP generation, whereby the membrane potential built by the proton pumps of respiratory chain complexes I, III, and IV is not matched by proton reentry through the mitochondrial F_1_F_0_ ATPase. In these conditions, ATP synthesis would be limited while mitochondrial membrane potential would increase, giving rise to increased ROS production and mitochondrial oxidative damage. It is known that mitochondrial ROS production is positively correlated with mitochondrial membrane potential (Turrens, 2003) and that oxidative stress causes mitochondrial fragmentation through activation of mitochondrial fission and inhibition of fusion (Iqbal & Hood, 2014). This proposed mechanism is consistent with prior *in vivo* ^31P-magnetic resonance spectroscopy (MRS) data demonstrating mitochondrial deficits and reduced ATP levels in presymptomatic DRPLA individuals (Lodi *et al*, 2000). Together, these findings underscore the notion that mitochondrial dysfunction and oxidative stress are early and critical drivers of DRPLA pathogenesis. Importantly, this phenotype occurs in the absence of heightened neuronal activity, as supported by MEA recordings showing preserved firing and network synchrony in early-stage DRPLA neurons, arguing against excitotoxic mechanisms and instead implicating a cell-autonomous dysregulation of mitochondrial homeostasis (Britti *et al*, 2018).

Morphological analyses revealed a progressive shift toward fragmented, rounded mitochondria; features typically associated with stress adaptation, impaired oxidative phosphorylation, and altered quality control (Fogo *et al*, 2024). While the molecular triggers of this hyperpolarized, fragmented state remain incompletely defined, it was accompanied by transcriptional signatures of oxidative stress, including downregulation of glutathione metabolism and suppression of mitochondrial thioredoxin reductase TXNRD2 (log FC -1.99). Although NQO1(log FC 1.49), a canonical NRF2 target, was induced, the overall antioxidant response was reduced, suggesting that DRPLA neurons mount a compensatory but incomplete redox defense in the face of elevated ROS burden. In the setting of elevated ROS production, consistent with early oxidative stress and redox imbalance. Although ATN1 inclusions and unfolded protein response (UPR) activation were not detected at baseline, the accumulation of autolysosomes suggests impaired or delayed cargo degradation. This may hinder the clearance of fragmented mitochondria, thereby sustaining mitochondrial dysfunction. Another possibility is that endoplasmic reticulum (ER) stress may contribute to this phenotype.

In addition to these early metabolic defects, our model captures both developmental and degenerative phenotypes of DRPLA. DRPLA patient-derived neurons exhibit a premature generation of CTIP2+ deep-layer fates, accompanied by a sustained reduction in SATB2+ upper-layer neuron generation. These observations suggest a shift in cortical fate, potentially reflecting accelerated deep-layer differentiation that prematurely exhausts the progenitor pool required for upper-layer neurogenesis. These fate shifts may be modulated by early mitochondrial and redox imbalance, as metabolic state and oxidative stress are known regulators of progenitor dynamics and fate transitions during corticogenesis (Chui *et al*, 2020; Khacho *et al*, 2019).

From a therapeutic perspective, we found that PB significantly improves neuronal survival and reduces oxidative stress, outperforming TUDCA alone and PB and TUDCA in combination. The precise mechanism of PB action remains incompletely defined; it likely acts through multiple pathways, including reduction of mitochondrial membrane potential, and enhancement of proteostasis via its chemical chaperone activity and HDAC inhibitory functions (Singh *et al*, 2025; Tanis *et al*, 2015). Sodium butyrate, a derivative of PB, has previously been shown to improve motor function and survival in a DRPLA mouse model by enhancing histone acetylation, further supporting PB as a therapeutic agent in DRPLA (Ying *et al*, 2006). Notably, both PB and TUDCA reduced survival in control cultures, suggesting that their beneficial effects may be selectively advantageous in the context of cellular stress.

Future studies should investigate how soluble ATN1 perturbs mitochondrial redox state and ROS production to further elucidate its role in DRPLA. Additionally, understanding the molecular underpinnings of SATB2+ neuronal vulnerability will be crucial for deciphering the selective degeneration of cortical neurons in DRPLA.

### Ethics approval and consent to participate

Written informed consent was obtained from all participants in accordance with the Declaration of Helsinki, permitting the use of biological material for research purposes only. The study was conducted with ethical approvals under the Institutional Review Board of Weill Cornell Medicine (IRB Protocol #1402014802).

## Materials and Methods

### Patient Sample Collection

Written informed consent was obtained from all participants in accordance with the Declaration of Helsinki, permitting the use of biological material for research purposes only. Four participants (three males and one female) were selected for their diagnosis of DRPLA associated with 66 or 67 CAG repeats in the ATN1 gene (EV Figure 1). Fibroblast primary cell lines were established from skin biopsies (3 mm) collected from DRPLA patients under a protocol approved by the Institutional Review Board of Weill Cornell Medicine (IRB Protocol #1402014802) for the purpose of reprogramming into pluripotent stem cells.

Two control human induced pluripotent stem cell (iPSC) lines derived from healthy individuals were used in this study. The first, designated CTRL1 (050877-01-MR), was derived from a 26-year-old male donor of European ancestry and obtained from the NY Stem Cell Foundation (NYSCF). CTRL2 was a gift from the University of Michigan. Both lines were reprogrammed from fibroblasts using the four reprogramming factors identified by Yamanaka (Takahashi *et al*, 2007). These lines tested negative for mycoplasma, retained normal karyotypes, and expressed canonical pluripotency markers.

### Derivation and Reprogramming of Patient-Derived Fibroblasts into Induced Pluripotent Stem Cells

Primary dermal fibroblasts were cultured using standard aseptic techniques in Dulbecco’s Modified Eagle Medium/Nutrient Mixture F-12 (DMEM/F12; ThermoFisher Scientific, Cat#10565018,), supplemented with 10–20% fetal bovine serum (FBS; ThermoFisher Scientific, Cat#10565018) and 1% penicillin–streptomycin (ThermoFisher Scientific, Cat# 15140122). Cells were maintained at 37 °C in 5% CO₂ and passaged at ∼80% confluence using TrypLE Express (ThermoFisher Scientific, Cat# 12604021).

Reprogramming of D66M, D67M1, D67M2, and D67F fibroblasts into human induced pluripotent stem cells (hiPSCs) was conducted by the Translational Research Core (CTRC) at the University of Utah using the CytoTune™ Sendai-OKSM viral vector system (infection ratio 5,5,3) (ThermoFisher Scientific, Cat# A16518), following established protocol. Fibroblasts were transduced with Sendai viral vectors encoding OCT4, SOX2, KLF4, and c-MYC at a multiplicity of infection (MOI) ratio of 5:5:3. After 24 hours of viral transduction, the culture medium was replaced daily. On day 7 post-transduction, cells were replated onto vitronectin-coated plates (ThermoFisher Scientific, A14700) and gradually transitioned to Essential 8 (E8) medium (ThermoFisher Scientific, A1517001). iPSC colonies with defined epithelial morphology emerged between days 18 and 21 and were manually picked and expanded in E8 medium. These colonies were expanded and cryopreserved at passage 5.

Pluripotency was confirmed by immunocytochemistry using antibodies against OCT4, SOX2, and NANOG, each applied at a dilution of 1:500. All iPSC lines were further validated for genomic stability (ThermoFisher Scientific, KaryoStat). Genetic identity was confirmed using short tandem repeat (STR) profiling. To ensure cell line integrity, all cultures were tested for the absence of mycoplasma contamination using e-Myco PLUS Mycoplasma PCR Detection Kit (iNtRON Biotechnology, Cat# 25237). All subsequent experiments were performed using early-passage iPSC lines (Passage 5 to P10) to minimize the risk of genetic drift and differentiation potential.

### Generation of cortical neural progenitor cells from induced pluripotent stem cells

iPSCs were seeded onto a Geltrex-coated plate (ThermoFisher Scientific, cat# A1413302) and maintained in StemFlex medium (ThermoFisher Scientific, cat# A3349401). Neural induction was initiated using the dual-SMAD inhibition protocol, following the manufacturer’s guidelines (STEMdiff™ SMADi Neural Induction Kit, StemCell Technologies, Cat# 08582).

In brief, iPSCs were dissociated using ReLeSR (StemCell Technologies, Cat#100-0483), centrifuged at 800x *g* for 3 minutes, resuspended in neural induction medium (NIM; StemCell Technologies, cat# 08582) supplemented with 10 μM Y-27632 ROCK inhibitor, and plated into Aggrewell 800 plates (StemCell Technologies, Cat. No. 08582). After 24 hours, the medium was replaced with fresh NIM without ROCK inhibitor and was changed daily thereafter. Embryoid bodies (EBs) formed within the AggreWell microwells and were transferred on day 5 to Geltrex-coated plates for adherent culture in NIM.

Neuroepithelial rosettes typically emerged between days 5 and 7 and were dissociated at day 12 following 1.5-hour incubation in STEMdiff Neural Rosette Select Reagent (StemCell Technologies, cat# 05832) and then replated onto Geltrex-coated plates in fresh NIM. Daily medium changes were continued until a homogenous population of neural progenitor cells (NPCs) emerged, at which point cultures were dissociated with Accutase and plated onto Geltrex-coated plates and transitioned to STEMdiff^TM^ neural progenitor medium (StemCell Technologies, cat# 05838) for maintenance and expansion. NPCs were characterized at passages 2 and 4 by immunocytochemistry for NPC markers NESTIN and SOX2, as well as dorsal forebrain identity markers PAX6, FOXP1, and FOXG1. Ventral identity was excluded by the absence of NKX2.1 expression (see Fig. 1). All control and DRPLA lines were differentiated and validated in parallel under identical conditions.

### Generation of cortical neurons from neural progenitor cells

Neural progenitor cells (NPCs) derived from control and DRPLA patient iPSCs were expanded up to passage 2, cryopreserved, and later thawed for future differentiation experiments. After thawing, NPCs were expanded for two additional passages and used at passage 4 for cortical neuronal differentiation and NPC studies.

For neuronal cultures, tissue culture plates were coated overnight at 37 °C with poly-L-ornithine (PLO; Sigma-Millipore, Cat# P4957), followed by three washes with sterile water and air-drying inside a biosafety cabinet. Plates were then coated with laminin (20 μg/mL) and fibronectin (5 μg/mL) in 1X PBS and incubated overnight at 37 °C. Cells were seeded in STEMdiff™ Forebrain Neuron Differentiation Medium (StemCell Technologies, Cat# 08600) and maintained for 7 days to initiate differentiation. Cultures were transitioned to BrainPhys™ Neuronal Medium (StemCell Technologies, Cat# 05793) supplemented with 20 ng/mL Brain-Derived Neurotrophic Factor (BDNF; PeproTech, Cat# 450-02), 20 ng/mL Glial cell line-Derived Neurotrophic Factor (GDNF; PeproTech, Cat# 450-10), 1 mM cyclic adenosine monophosphate (cAMP; StemCell Technologies, Cat# 73886), and 1 μg/mL laminin (mouse; Corning, Cat# 354232). Medium was refreshed every other day using a 50% medium exchange strategy to preserve endogenous trophic support while maintaining nutrient availability. All DRPLA and control NPC lines were differentiated in parallel under identical conditions. Neuronal identity and purity were confirmed by immunocytochemistry for MAP2 and βIII-Tubulin (TUJ1). Cortical layer specification was assessed using CTIP2 and SATB2.

### Multielectrode Array (MEA)

To evaluate spontaneous extracellular electrophysiological activity, cortical neurons were plated on 48-well Cytoview MEA plates (Axion Biosystems, Cat# M768-tMEA-48W-5) and recorded using the Maestro MEA system (Axion Biosystems, Cat# M768-tMEA-48W-5). Prior to cell seeding, MEA plates were pre-coated with PLO and incubated at 37°C for 18 hours, followed by three washes with sterile water. Plates were then air-dried under sterile conditions in a laminar flow hood. A second coating of Laminin (20 μg/ml) and fibronectin (5 μg/ml) (Sigma Millipore, Cat# F4759), diluted in 1X PBS, was added to each well and incubated at 37 °C for an additional 18 hours to promote neuronal adhesion and maturation.

NPCs were seeded at a density of 50,000 cells per well into 16 wells per line. On day 3 post-seeding, primary mouse astrocytes (20,000 cells per well) were added to the wells to support neuronal connectivity and synaptogenesis. Following astrocytes addition, a proliferation inhibitor, 5-Fluoro-2′-deoxyuridine (Sigma-Aldrich, Cat#F0503), was added to prevent astrocyte overgrowth.

### MEA recordings

Spontaneous extracellular neuronal activity was recorded every other day for 5 minutes. During recordings, MEA plates were kept at 37°C with 5% CO₂ for the duration of the recordings to preserve physiological conditions. Recordings were processed and analyzed using the AxIs Navigator Neural Metrics tool (Axion Biosystems). The system was set to measure AxIS Raw, AxIS Spike, and Network Burst List. Spike files were processed with Neural Metrics software. Analysis Parameters: active electrode criterion 1 spike/minute, with coincident artifacts removed—Burst Parameters: ISI threshold minimum numbers of spikes 5, max ISI 100 ms. Network Burst Parameters: Envelope threshold factor 1.25, minimum IBI 100 ms, minimum electrodes 25%, burst inclusion 75%. Synchrony Parameters: Normalized cross-correlogram, synchrony window 20 ms, Synchrony Index. Average Network Burst Parameters: events 1-N, window 0-500 ms, bin size 1 ms. To refine burst detection and reduce noise artifacts, an inter-spike interval (ISI) threshold of 300 milliseconds was applied during spike train analysis.

### Molecular Cloning and Lentivirus Generation

Plasmid constructs Synapsin-EGFP (Addgene, Cat#114215) and FUW-mCherry-GFP-LC3 (Addgene, Cat#110060) were obtained from Addgene under a Material Transfer Agreement (MTA). Lentiviral particles were produced in HEK293T cells cultured in standard 10 cm dishes. Cells were seeded to reach 80–90% confluency at the time of transfection.

For each dish, cells were co-transfected with 7.5 µg of the transfer plasmids (either Synapsin-EGFP or FUW-mCherry-GFP-LC3), 4.5 µg of the packaging plasmid psPAX2 (Addgene, Cat#12260), 1.5 µg of the envelope plasmid pMD2.G (Addgene, Cat#12259), using polyethylenimine (PEI; Sigma-Aldrich, Cat# 40872) as the transfection reagent. Plasmids were mixed in 600 µl of Opti-MEM reduced serum media (ThermoFisher Scientific, Cat# 31985062) with 48 µl of PEI (1 mg/ml), incubated briefly at room temperature, and added dropwise to the cells.

After 18 hours of post-transfection, the medium was replaced by fresh Dulbecco’s minimal essential medium (DMEM) (ThermoFisher Scientific, Cat# 11330032), supplemented with 10% FBS and 1% penicillin-streptomycin. Viral supernatant was collected at 48- and 72-hours post-transfection, centrifuged at 1,200 *g* for 5 minutes at 4°C to remove cell debris, and filtered through 0.45 µM polyethersulfone (PES) syringe filters.

For viral concentration, the filtered supernatant was mixed with Lenti-X Concentrator (Takara Biosciences, Cat# 631231) at ratio of 1:3 (concentrator: supernatant), gently inverted to mix, and incubated at 4°C for 30 min to overnight. The mixture was centrifuged at 1,500 x g for 45 minutes at 4°C. The resulting viral pellet was resuspended in either 1/10 or 1/100 of the original volume using complete DMEM stored at -80 °C until use.

### Neuronal viability tracking by longitudinal EGFP imaging

To assess long-term neuronal survival, iPSC-derived cortical neurons were labeled with a lentiviral vector expressing EGFP under the human Synapsin promoter (pHR-hSyn-EGFP, Addgene #114215). NPCs were seeded at a density of 20,000 cells per well on PLO and laminin-coated 96-well plates. At day 1 of differentiation, NPCs were transduced with the EGFP expressing lentiviral vector at a total of 0.5X10^6^ transfection units per well without polybrene. Automated imaging was performed every two days starting at D13 of differentiation and continued until day 80. Imaging was conducted using the ImageXpress Pico High-Content Imaging System (Molecular Devices) at 10X magnification.

Acquired images were processed in Fiji. Background fluorescence was subtracted, and image contrast was adjusted uniformly across datasets. Neuronal cell bodies were identified by applying Gaussian blur, followed by intensity thresholding and size-based particle exclusion. The number of EGFP-positive neurons per field was quantified at each time point. To account for plating variations, cell counts were normalized to the initial timepoint D13, setting baseline survival to 100%. Neuronal survival was plotted over time to compare DRPLA and control lines.

### Reactive Oxygen Species (ROS) Detection using DHE

Cytosolic ROS levels were measured using the cell-permeable superoxide indicator dihydroethidium (DHE) (Cat no. D11347; ThermoFisher Scientific). Upon oxidation by superoxide, DHE is converted to 2-hydroxyethidium and other oxidation products, which intercalate with DNA and emit red fluorescence. Neurons were incubated with 5µM DHE diluted in Isosmotic Artificial Cerebrospinal Fluid or Intracellular Artificial Cerebrospinal Fluid (I-ACSF) for 30 minutes at 37°C. After incubation, cells were gently washed and replaced with fresh medium before confocal acquisition. Imaging was performed using an Olympus confocal microscope under live-cell conditions using a 20x objective. ROS quantification was performed by subtracting baseline background intensity from integrated DHE fluorescence using ImageJ, with images taken per neuronal soma.

### Detection of Mitochondrial ROS (mtROS) using MitoSOX Red

Mitochondrial ROS (mtROS) levels were assessed using MitoSOX Red (ThermoFisher Scientific, Cat# M36008), a mitochondrial-targeted fluorogenic dye selective superoxide indicator. A stock solution was prepared and diluted in Hank’s Balanced Salt Solution (HBSS) (ThermoFisher Scientific, Cat# 88284) to a final working concentration of 500 nM, following the manufacturer’s instructions. Live cells were incubated with the MitoSOX solution for 30 minutes at 37 °C, followed by washing with HBSS and the addition of fresh culture medium before imaging. Confocal images were acquired using Olympus under identical acquisition settings across samples. Quantification of MitoSOX intensity was performed on individual EGFP-labeled neurons using ROI-based analysis, and averaged fluorescence per neuron was used as the readout for mtROS levels.

### Mitochondria Assays

Mitochondria membrane potential was evaluated with Tetramethylrhodamine, Methyl Ester, and Perchlorate (TMRM) (ThermoFisher Scientific, Cat# T668) at D10 and D20 of differentiation. Live cells were incubated with TMRM at a final concentration of 15 nM for 45 minutes in BrainPhys medium. Live imaging was performed on an ImageXpress Pico microscope (Molecular Devices) at a 40x objective (pixel size = 0.1725 μM). Single-cell segmentation was performed based on nuclear morphology and cytoplasmic boundaries. iPSCs-derived neurons were identified and filtered using morphological criteria. For each neuron, mean TMRM fluorescence intensity was quantified within the EGFP+ area after background subtraction. Individual mitochondria were segmented using intensity thresholds and skeletonized to measure mitochondria size, shape, and polarization. Measurements from individual mitochondria were averaged to generate per-cell values. To validate TMRM dynamic responsiveness, cultures were treated with the mitochondrial uncoupler carbonyl cyanide-p-trifluoromethoxy phenylhydrazone (FCCP;100 μM final concentration) (Millipore Sigma, Cat# C2920-10MG) followed by live imaging to confirm TMRM signal dissipation.

Mitochondrial morphology was additionally assessed using MitoTracker Red CMXRos (ThermoFisher Scientific, Cat# M7512). Live cells were incubated with MitoTracker (200 nM final concentration for 20 minutes at 37 °C in serum-free media, followed by live imaging with Olympus confocal microscopy.

### Immunocytochemistry

Cells were fixed with 4% paraformaldehyde (PFA) for 10 minutes at room temperature, followed by three washes in phosphate-buffered saline (PBS). Permeabilization was performed using 0.3% Triton X-100 in 1% normal donkey serum (NDS) for 20 minutes. Cells were then incubated overnight at 4 °C with primary antibodies diluted in blocking buffer (1% NDS in PBS with 0.1% Triton X-100). The following primary antibodies were used: rabbit anti-ATN1 (1:500; Millipore Sigma, HPA031619), mouse anti-polyglutamine (1:1000; Millipore Sigma, MAB1574), mouse anti-TUJ1 (1:1000; BioLegend, 801201), chicken anti-MAP2 (1:1000; ThermoFisher, PA1-10005), rat anti-CTIP2 (1:1000; Abcam, ab167575), rabbit anti-TBR1 (1:1000; Abcam, ab31940), goat anti-SOX1 (1:1000), rabbit anti-PAX6 (1:1000; MBL, PD022), rabbit anti-FOXG1 (1:300; Abcam, ab18259), chicken anti-Nestin (1:1000; Novus, NB100-1604), rabbit anti-cleaved Caspase-3 (1:500; Cell Signaling, 9661S), guinea pig anti-FOXP1 (1:16,000), rabbit anti-NKX2.1 (1:500; Abcam, ab76013), mouse anti-SATB2 (1:100; Abcam, ab51502), rabbit anti-phospho-Histone H3 (PHH3; 1:500; Cell Signaling), rabbit anti-MYT1L (1:1000; Millipore Sigma, ABE2915), and rabbit anti-Ki67 (1:2000; ThermoFisher, MA5-14520). After washing, species-specific Alexa Fluor-conjugated secondary antibodies (ThermoFisher) were applied at 1:1000 dilution for 1 hour at room temperature. Nuclei were counterstained with DAPI and mounted using ProLong™ Glass Antifade Mountant (ThermoFisher Scientific, Cat# P36980).

### Pharmacological inhibition of proteasome and autophagy pathways

Experiments inhibiting proteosome and autophagy pathways were carried out using inhibitor concentrations and timing as previously described by others (Han *et al*, 2009; Yamamoto *et al*, 1998). To inhibit proteasomal degradation, cells were treated with MG132 (5 μM; Selleckchem, Cat #S2619) for 18 hours. For autophagy inhibition, Bafilomycin A1 (BAF; 100 nM; Sigma-Aldrich, Cat# SML1661-.1ML) was dissolved in dimethyl sulfoxide DMSO (ThermoFisher Scientific, Cat# D12345) and added to the culture medium for 18 hours prior to imaging. All vehicle controls received an equivalent volume of DMSO (<0.1% final concentration). Treatments were performed in triplicate, and culture conditions were kept constant throughout all experimental groups.

### Fluorescence-Activated Cell Sorting (FACS) of neurons

Neurons were transduced with pSyn-EGFP lentiviral vector on day 1 of differentiation and harvested on day 20 for FACS-based isolation of live neurons. Cultures were rinsed with 1x PBS and incubated with Accutase (StemCell Technologies, Cat# 07920) at 37 °C for 3 minutes to dissociate cells. The resulting cell suspension was collected in DMEM/F-12 medium, centrifuged at 300x *g* for 5 minutes at 4 °C, and resuspended in HBSS supplemented with 5% FBS and DNase I (0.1 mg/mL).

Cell viability and neuronal identity were assessed with DAPI and GFP fluorescence. Dead cells were identified as DAPI-positive, while live neurons were defined as DAPI-negative and GFP-positive (DAPI-/GFP+). Gates strategy included exclusion of debris and doublets based on forward and side scatter. Gates were established using untransduced and pSyn-EGFP-transduced neural cultures as negative and positive controls, respectively. DAPI-/EGFP+ neurons were collected for downstream RNA sequencing. Flow cytometry was performed using (MA900 Cell Sorter, BD Biosciences) operated at a medium flow rate with a (100 μm) nozzle.

### RNA extraction

Total RNA was extracted and isolated with QiaZol reagent (Qiagen, Cat # 79036) and RNeasy mini kit (Qiagen, Cat #74104) following the manufacturer’s instructions. RNA concentration and integrity were assessed using. All samples used for bulk RNA seq had a RIN > 8.8.

### RNA sequencing

RNA samples were sent to the Weill Cornell Medicine Genomics Core for library preparation and sequencing. Sample libraries were generated using (NEB Ultra II Directional RNA Library Prep (plus Poly A isolation module). RNA kit and sequenced with paired-end 150 base pair reads on one lane using Illumina HiSeq 2500, with 40 million reads.

### Data Normalization and Quality Control

Raw RNA-seq count data were normalized using the EDASeq package (Bioconductor, v2.38.0) to correct for within- and between-lane technical effects. Quantile normalization was subsequently performed using the preprocessCore package (v1.68.0), and normalized counts were log₂-transformed prior to downstream analysis. All data processing and statistical analyses were performed in R (v4.4.1) using RStudio (v2023.06.1).

To evaluate global transcriptional variance and sample clustering, principal component analysis (PCA) was performed using the base R prcomp function. PCA plots were generated using ggplot2 (v3.5.2) to visualize genotype-specific clustering and assess potential outliers.

### Differential Expression Analysis

Differential gene expression between DRPLA and control iPSC-derived neurons was analyzed using the limma package (v3.60.4), with voom normalization to estimate the mean–variance relationship and apply precision weights, followed by empirical Bayes moderation. Genes were considered significantly differentially expressed if they exhibited a Benjamini–Hochberg adjusted p-value < 0.05 and an absolute log₂ fold-change > 1. Top-ranked genes were visualized using volcano plots generated in ggplot2, with annotated fold-change and adjusted p-value thresholds.

### Pathway Enrichment Analysis

Pathway enrichment analysis was performed using Ingenuity Pathway Analysis (IPA; QIAGEN). All enriched pathways were derived from the IPA knowledge base. Enrichment significance was determined using the IPA default settings, including Benjamini– Hochberg correction for multiple comparisons. Results were visualized as stacked bar plots using ggplot2 (v3.5.2), and summary figures were prepared using GraphPad Prism (v10).

### Western Blots

NPC or neurons were rinsed with cold 1x Phosphate-buffered saline (DPBS, 1X), Dulbecco’s formula, without calcium, without magnesium (137 mM Sodium chloride, 8.1 mM disodium phosphate, 1.47 mM monopotassium phosphate, and 2.7 mM potassium chloride, pH 7.4). Cells were lysed on ice for 30 minutes with RIPA buffer (1% Triton X-100, 1% sodium deoxycholate, 1% SDS, 0.1%, 150 mM NaCl, 10 mM Tris-HCl, pH 7.4) supplemented with protease inhibitors. Lysates were clarified by centrifugation at 10,000x *<u>g</u>* for 10 minutes at 4°C.

Protein concentrations were determined using Pierce™ BCA Protein Assay Kit (ThermoFisher Scientific, Cat# 23225) and normalized using the corresponding lysis buffer. Lysates were mixed with 4x sample buffer containing 50 mM DTT and heated at 37 °C for 10 minutes. Samples were resolved on either NuPAGE™ 4-12% Bis-Tris Plus Mini Protein Gels (ThermoFisher Scientific, Cat# NW04125BOX) in MES SDS running buffer or on conventional 4% SDS-PAGE gels in Tris-Glycine-SDS buffer. Proteins were transferred onto PVDF membrane (Bio-Rad Laboratories, Cat# 1620177) in a transfer buffer (25 mM Tris, 192 mM glycine, pH 8.3,10% Methanol) at 15V, 4°C overnight.

Membranes were blocked with 1% BSA in TBST (50 mM Tris-Cl, pH 7.5; 150 mM NaCl, 0.1% Tween 20) for 1 hour at room temperature and incubated overnight at 4°C with primary antibodies diluted in 1% BSA in TBST, blotted with anti-ATN1 (rabbit polyclonal, 1:500; Millipore Sigma, HPA031619-100), and anti-a-tubulin (mouse monoclonal; 1:2000, Proteintec, 66031-1-Ig). Membranes were washed 3 times with 1X TBST for 10 minutes each, followed by a final rinse with 1X TBS. Detection was performed using Immobilon Crescendo Western HRP substrate (Millipore Sigma, Cat# WBLUR0500), and signals were imaged using the Odyssey Fc Imaging system (LI-COR Biosciences). Band intensities were quantified using ImageJ software.

### Pharmacological Treatments

Phenylbutyrate (PB; Selleckchem, Cat# S4075) and tauroursodeoxycholic acid (TUDCA; Sigma-Aldrich, Cat# T0266) were dissolved in sterile water and applied to neuronal cultures either as monotherapies or in combination across a range of concentrations. PB was used at final concentrations of 1000 µM, 500 µM, and 100 µM, while TUDCA was used at 100 µM, 50 µM, and 25 µM. Combination regimens included (1000 µM PB + 100 µM TUDCA), (1000 µM PB + 50 µM TUDCA), (500 µM PB + 50 µM TUDCA), (500 µM PB + 25 µM TUDCA), and (100 µM PB + 10 µM TUDCA). Treatments were initiated on day 1 of neuronal differentiation and applied daily in neuronal differentiation medium for the first 7 days. From day 8 onward, treatments were continued every other day in BrainPhys medium for the duration of the experiment. To control for vehicle effects, equivalent volumes of sterile water were added to untreated control wells under identical medium change schedules.

### Image acquisition

Confocal fluorescence imaging was performed using an Olympus Fluoview FV1000 confocal microscope, operated with FV10-ASW software (v4.2.2.1). All images were processed using FluoView for initial export and visualization. Brightfield images were captured using an Olympus EP50 camera mounted on a standard inverted microscope. For high content imaging applications, including TMRM fluorescence and survival assays, images were acquired using a Pico automated imaging platform and analyzed using MetaMorph software (MetaSeries version 2.0.0.0). Where applicable, linear brightness and contrast adjustments were applied uniformly across conditions using Adobe Photoshop, strictly for figure presentation. No nonlinear processing, selective enhancement, or alterations were made to the raw data used for quantification.

### Data analysis and statistical methods

Statistical analyses were performed using GraphPad Prism version 10 (GraphPad Software). As described in the patient sample sections, experiments were conducted using two control iPSC lines and four DRPLA iPSC lines. Each line was subjected to three independent differentiations (n=3 per line), representing biological replicates. Within each differentiation, multiple technical replicates (e.g., neurons or fields of view) were averaged to yield a single value per replicate. In some experiments, data were analyzed at the genotype level by grouping iPSC lines. For these comparisons, the mean value from each of the three differentiations per iPSC line was used, and data were grouped across the two control lines (total n=6) and the four DRPLA lines (total n=12) for statistical testing. Comparisons between the two groups were performed using unpaired two-tailed Student’s *t*-tests, and statistical significance was defined as p < 0.05. Exact p-values and statistical tests used are reported in the figure legends.

## Declarations

There are no declarations to disclose.

## Acknowledgements.

We thank the Ross laboratory for their thoughtful feedback. We also acknowledge Michael Harrison and Shawn Singh for technical assistance. This work was supported by the Center for Neurogenetics and gifts from the Feil Family Foundation and the Compton family.

## Author’s contribution

CRediT contributor roles: Conceptualization: M.E.R. Data curation: N.A.N, C.K. Formal analysis: N.A.N, C.K, C.W, S.S and C.R. Funding acquisition: M.E.R. Investigation; N.A.N, C.K, C.W. Methodology: S.L, N.A.N, C.K, G.M, C.A.P and M.E.R. Project administration: N.A.N, C.A.P and M.E.R. Resources: S.L, G.M, S.S and C.R. Software: C.K, G.M, S.S and C.R. Supervision: G.M, C.A.P and M.E.R. Validation: N.A.N. Visualization: N.A.N and C.A.P. Writing - original draft: N.A.N, C.A.P and M.E.R. Writing - review & editing: N.A.N, C.K, G.M, C.A.P and M.E.R.

## Expanded View Figures and Figure Legends

**EV Figure 1.**
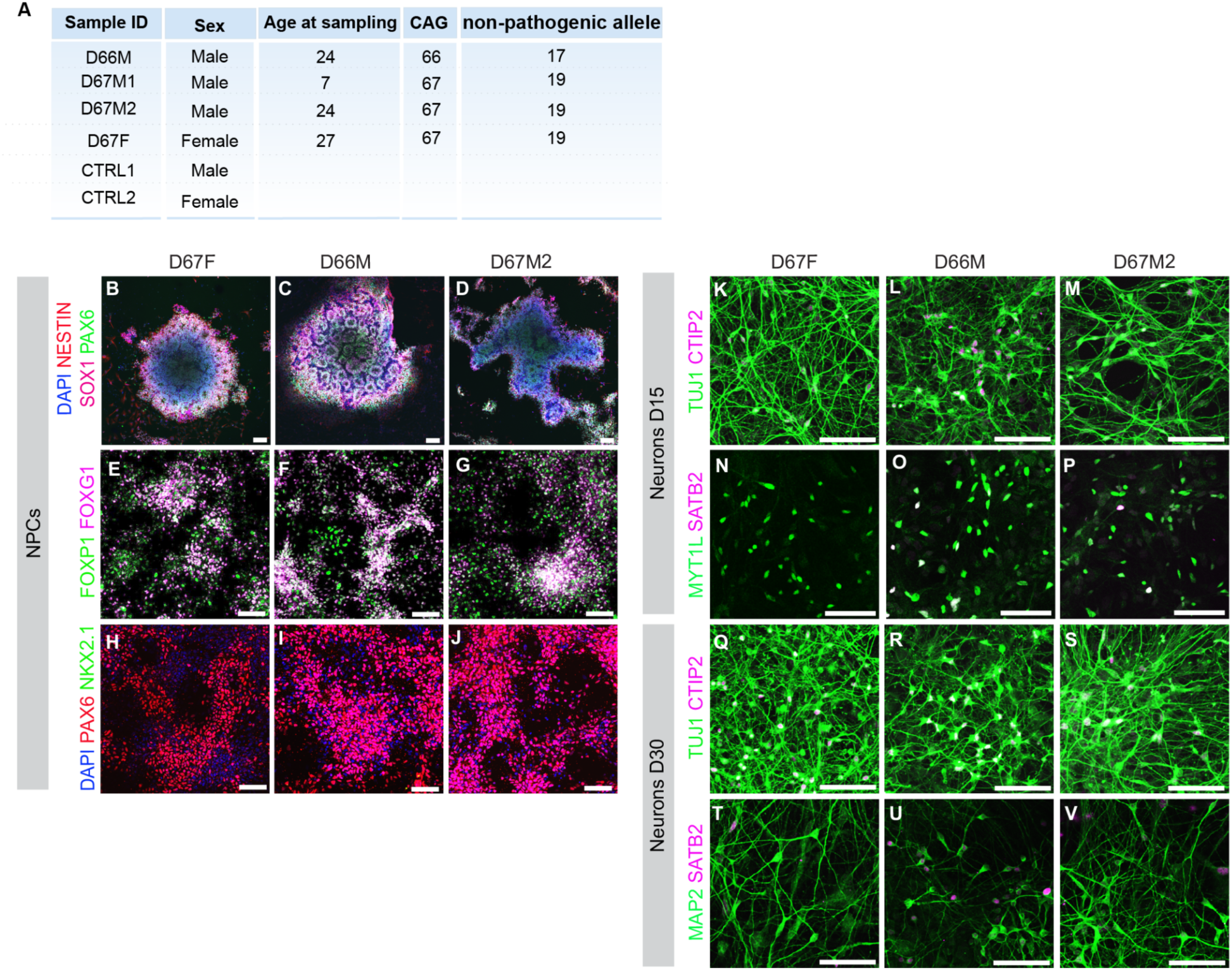
DRPLA patient-derived neurons differentiate appropriately towards cortical fate (related to Figure 1). (A) Table detailing DRPLA patient-derived ID numbers, sex, age at sampling, and number of CAG repeats in pathogenic and non-pathogenic alleles. (B-D) Immunostaining for NESTIN, SOX1, and PAX6 in neural rosettes selected from DRPLA patient-derived lines D67F, D66M and D67M2. (E-G) Immunostaining for FOXP1 and FOXG1 in neural rosettes selected from DRPLA patient-derived lines D67F, D66M and D67M2. (H-J) Immunostaining for PAX6 and NKX2.1 in neural rosettes selected from DRPLA patient-derived lines D67F, D66M, and D67M2. (K-P) Immunostaining for TUJ1 and CTIP2, and MYT1L and SATB2 in DRPLA patient-derived neurons D67F, D66M, and D67M2 at day 15 of differentiation. (Q-V) Immunostaining for TUJ1 and CTIP2, and MAP2 and SATB2 in DRPLA patient-derived neurons D67F, D66M, and D67M2 at day 30 of differentiation. Scale bars (B-C) 100 μM, (E-J) 50 μM, (K-V) 50 μM.

**EV Figure 2.**
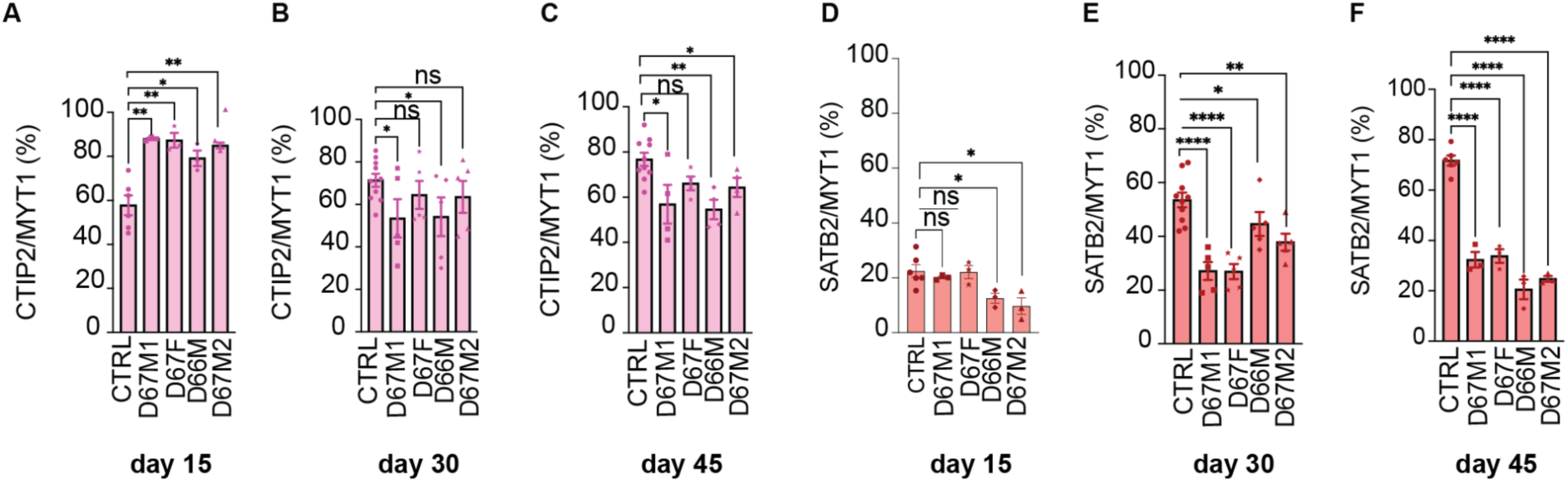
Premature differentiation towards deep layer neuronal fate in DRPLA patient-derived neurons (related to Figure 2). (A-C) Quantification of the proportion of MYT1L+ neurons expressing CTIP2 in control and DRPLA patient-derived neurons at days 15, 30, and 45. (D-F) Quantification of the proportion of MYT1L+ neurons expressing SATB2 in control and individual DRPLA patient-derived neurons at days 15, 30, and 45. Data represent mean ± SEM. N=2 control and 4 patient-derived biological replicates, 3 technical replicates. Statistical analysis was performed using a two-tailed Student’s t-test. p= **0.0025 (D67M1), **0.0038 (D67F), *0.0184 (D66M), **0.0047 (D67M2) (A); *0.0345 (D67M1),p=*0.0429 (D66M) (B); *0.0140 (D67M1), **0.0013 (D66M), *0.0381 (D67F2)(C); *0.0269 (D66M), *0.9129 (D67F2) (D);****<0.0001 (D67M1), ****<0.0001 (D67F), *0.0169 (D66M), **0.0018 (D67M2) (E);****<0.0001 (D67M1), ****<0.0001 (D67F), ****<0.0001 (D66M), ****<0.0001 (D67M2) (F).

**EV Figure 3.**
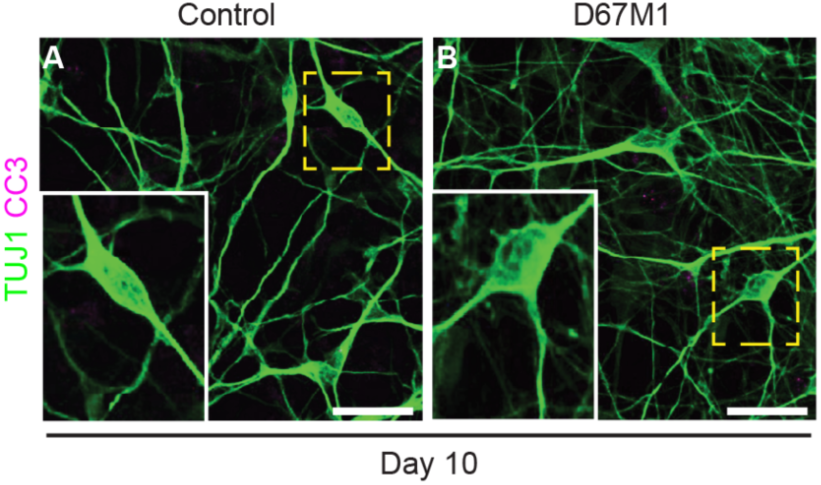
Apoptosis is not observed in DRPLA patient-derived neurons at day 10 (related to Figure 3). (A-B) Immunostaining of cleaved Caspase-3 (CC3) and TUJ1 in control and DRPLA (D67M1) patient-derived neurons at day 10 of differentiation. Scale bars: 20 μM.

**EV Figure 4.**
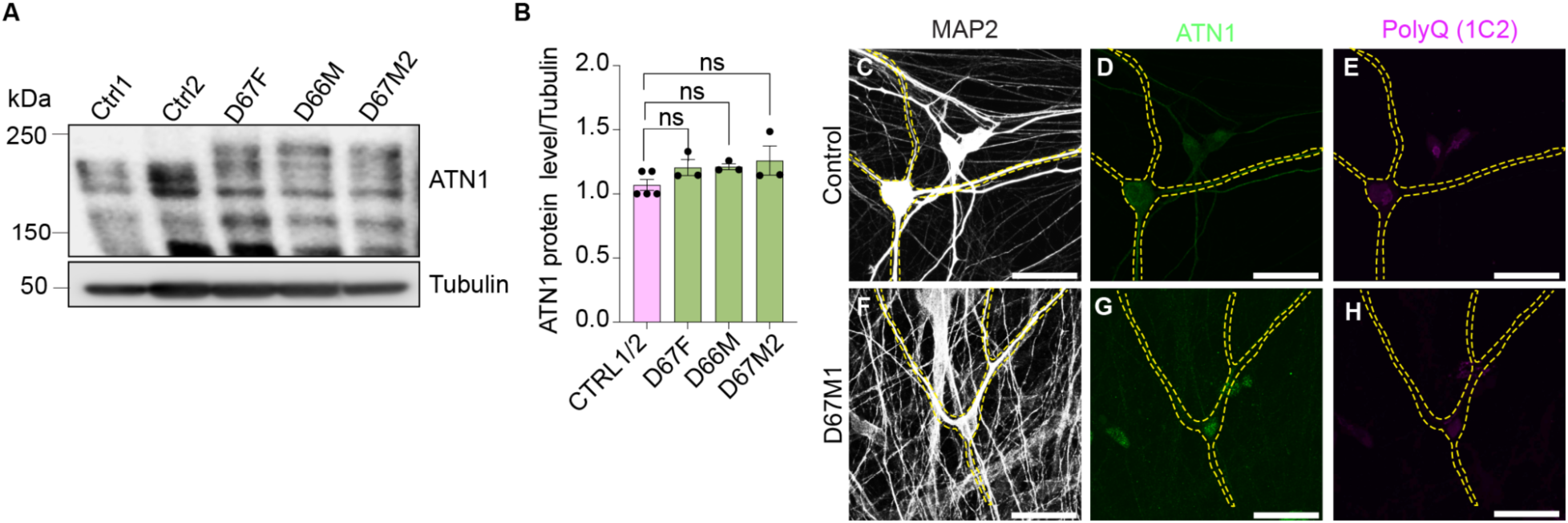
Polyglutamine inclusions are absent and ATN1 expression is unaltered in DRPLA patient-derived neurons after prolonged culture (related to Figure 4). (A) Western blot analysis of ATN1 expression in control and DRPLA patient-derived neurons at day 20 of differentiation. (B) Quantification of western blot intensity compared to Tubulin in control and DRPLA patient-derived neurons at day 20 of differentiation. (C-H) Immunostaining of MAP2, ATN1, and PolyQ (1C2) in control and DRPLA patient-derived neurons at day 70. Data represent mean ± SEM. N=2 control and 4 patient-derived biological replicates, three technical replicates. Statistical analysis was performed using a two-tailed Student’s t-test. p values (B) 0.1131 (D67F), 0.0563 (D66M), 0.1093 (D67M2). Scale bars: 20 μM.

**EV Figure 5.**
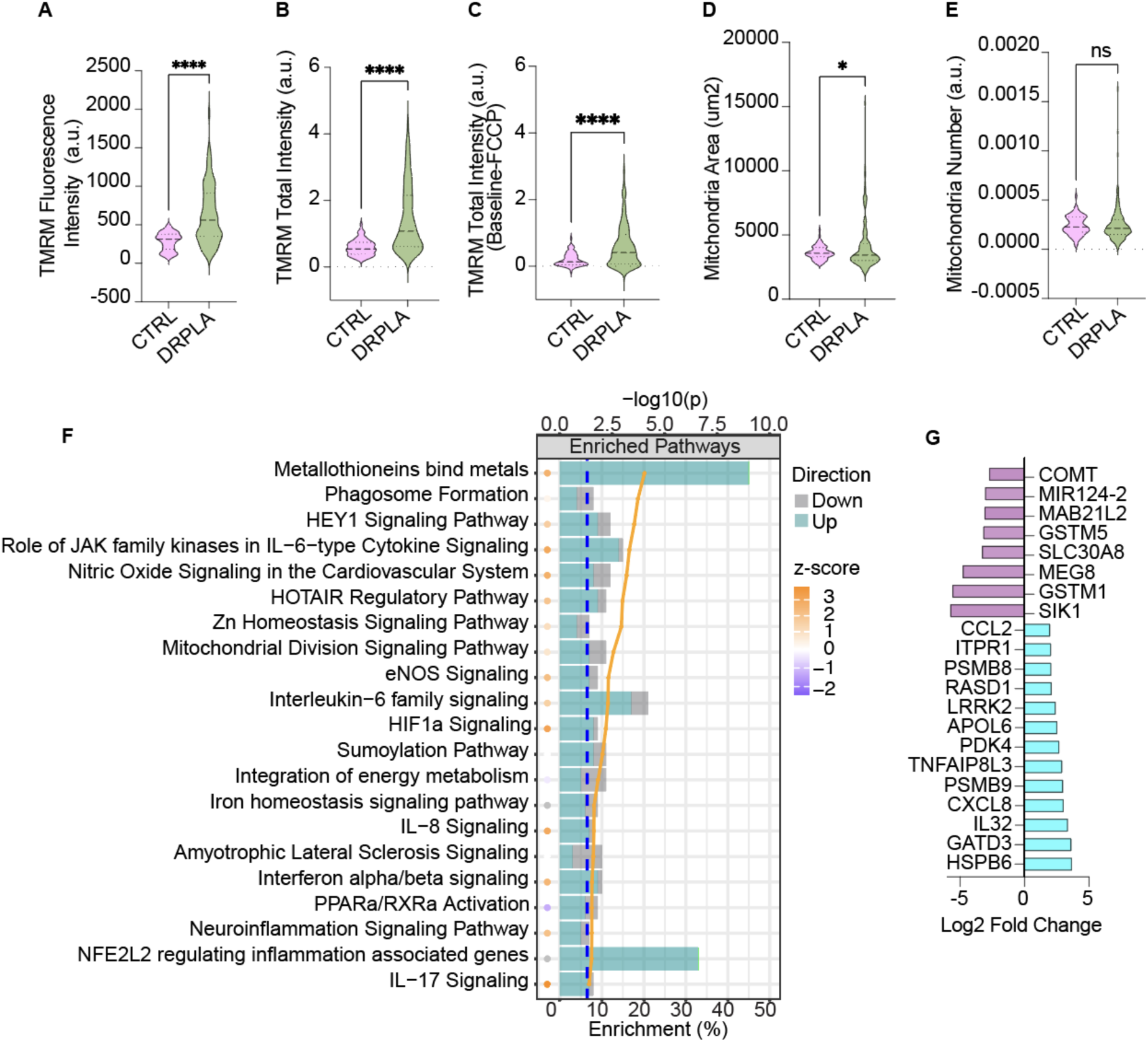
Increased oxidative stress is detected in DRPLA patient-derived neurons (related to Figure 5). (A-E) Quantification of TMRM fluorescence intensity, total intensity, total intensity -FCCP, and mitochondrial area and number in control and DRPLA patient-derived neurons at day 10. (F) Mitochondrial and oxidative stress-related GO pathways were identified as misregulated in DRPLA patient-derived neurons through RNA Seq. (G) Significantly misregulated genes in DRPLA patient-derived neurons identified by RNA Seq. Data represent mean ± SEM. N=2 control and 4 patient-derived biological replicates, 3 technical replicates. Statistical analysis was performed using two-tailed Student’s t-test. p= ****<0.0001 (A); ****<0.0001 (B); ****<0.0001 (C); *0.0442 (D).

**EV Figure 6.**
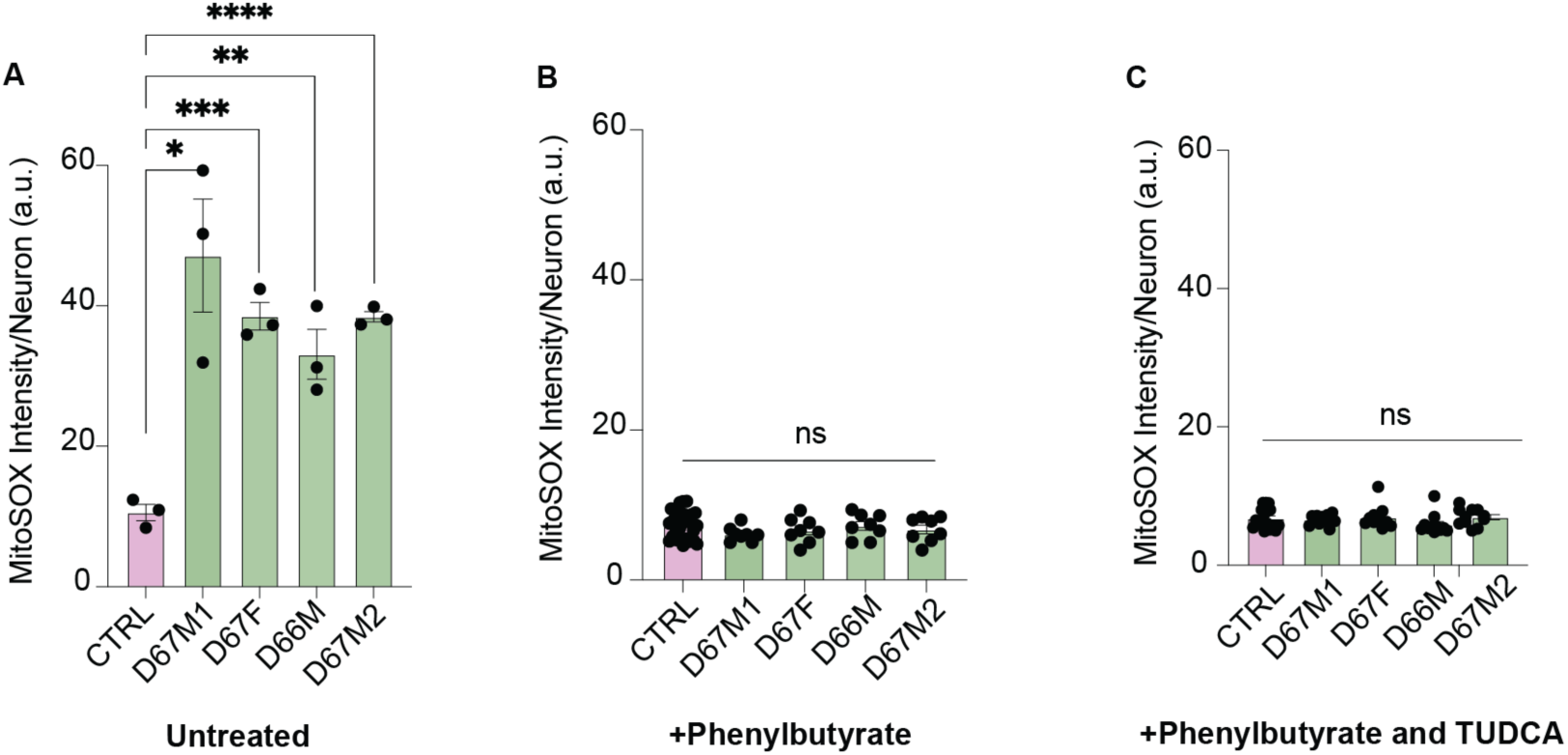
PB, TUDCA, and combination treatments reduce oxidative stress in DRPLA patient-derived neurons (related to Figure 6). (A) MitoSOX Red intensity in untreated control and DRPLA patient-derived neurons at day 20. (B) Mitosox intensity in control and DRPLA patient-derived neurons treated with phenylbutyrate at day 20. (C) Mitosox intensity in control and DRPLA patient-derived neurons treated with phenylbutyrate and TUDCA at day 20. Data are presented as mean ± SEM; each dot represents one field of view. N= 2 control and 4 patient-derived biological replicates, three technical replicates. Statistical analysis was performed using a two-tailed Student’s t-test. p=*0.0108 (D67M1), ***0.0003(D67F), **0.0039(D66M), ****<0.0001 (D67M2) (A).

**EV Figure 7:**
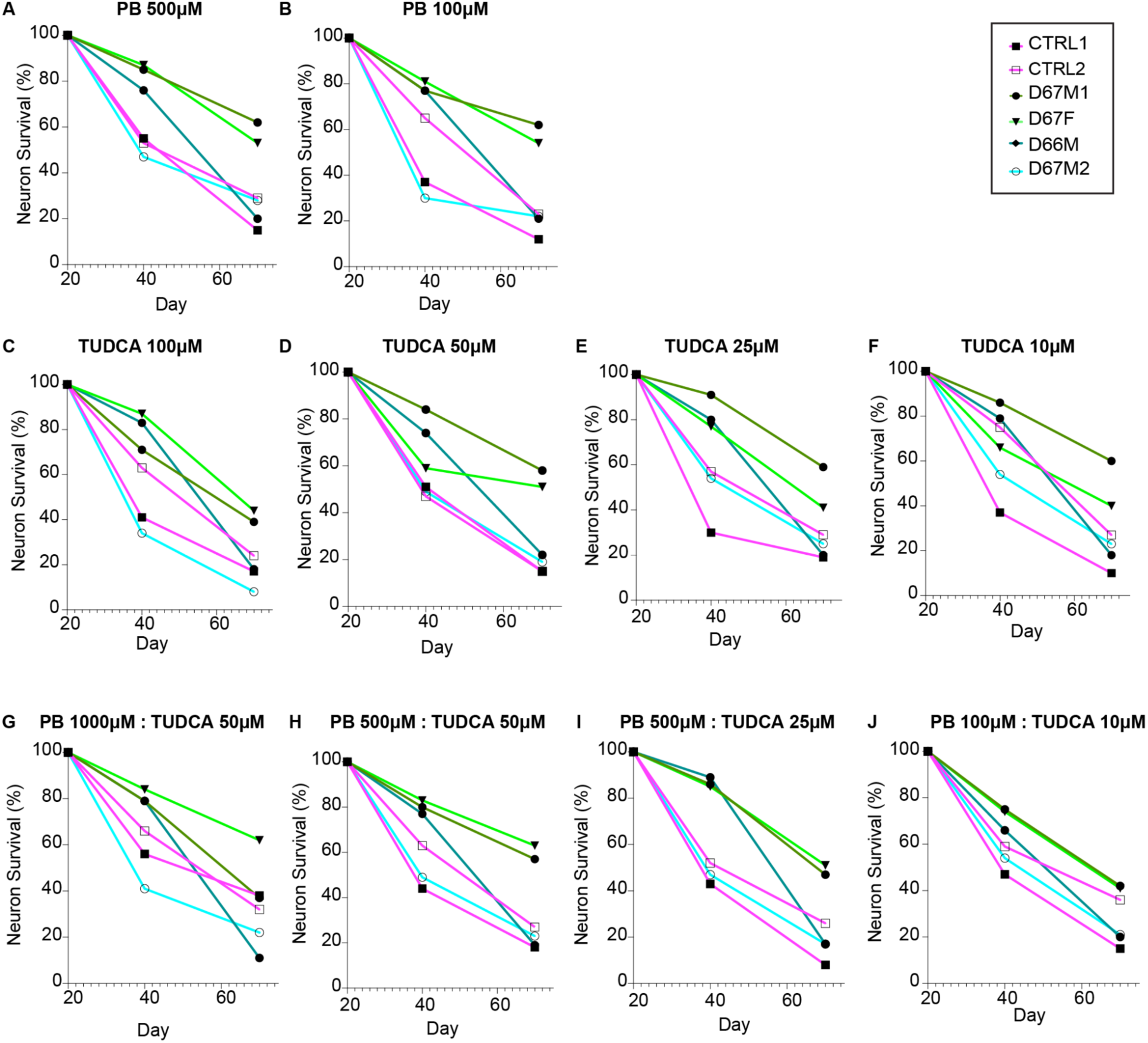
Longitudinal analysis of neuronal survival. DRPLA iPSC-derived neurons treated with suboptimal concentrations of PB, TUDCA, and PB-TUDCA combinations (related to Figure 7). **(A-J)** Dose-dependence analysis of neuronal survival from day 20 to day 70 of differentiation in DRPLA patient-derived (D67M1, D67M2, D67F, and D66M) and control (CTRL1, CTRL2) cortical neurons treated with phenylbutyrate (PB), TUDCA, or PB-TUDCA combinations. Survival was quantified as the percentage of EGFP+ neurons normalized to day 20 levels. Treatment was initiated at day 1 and maintained until day 70. Each line represents the mean survival of three independent differentiations per line. Survival plots using optimal concentrations of PB and TUDCA are shown in Figure 7.

